# Endemic koloa maoli (Hawaiian Duck, *Anas wyvilliana*) shows preferential social associations, but not based on plumage or genetic relatedness

**DOI:** 10.64898/2026.01.29.702521

**Authors:** P. Keerthipriya, C. P. Malachowski, B. D. Dugger, K. J. Uyehara, A. Engilis, P. Lavretsky, C.P. Wells

## Abstract

Island endemic birds are under greater threat than their mainland counterparts. Sedentary living and historically reduced predation may affect island bird sociality and inform their conservation and management. However, detailed studies on their sociality are uncommon. The federally- endangered koloa maoli (*Anas wyvilliana,* Hawaiian duck, or koloa) is primarily threatened by hybridization with feral mallards and avian botulism outbreaks. We used capture-mark-recapture and genetic data on koloa on the island of Kaua‘i, a stronghold of remaining koloa, to construct social networks and examine their associations (inferred from co-occurrence in traps) and spatial genetic structure. Information on associations might shed light on preferences for or against mallards and hybrids, and inform planned translocation efforts. Microgeographic spatial genetic structuring where relatedness among individuals scales with geographic distance is a potential liability for maintaining koloa genetic diversity, and would particularly be detrimental during highly localized outbreaks of botulism that could result in the removal of entire lineages. While we found persistent social associations among adult koloa, they were not apparently influenced by plumage traits or body mass, suggesting a lack of social preference for mallard-like individuals. Importantly, we found no spatial patterns of relatedness within the largest refuge. Therefore, botulism outbreaks remain a demographic threat but are unlikely to remove correlated genetic diversity. There were no sex differences in spatial genetic structure and both sexes moved within a refuge. The lack of spatial genetic structure and the presence of many unrelated conspecifics may enable koloa to limit inbreeding and retain genetic diversity without sex-biased dispersal. In the context of future translocations, our results suggest that translocating koloa captured in the same trap together will reduce disruption of preferred associations while also retaining genetic diversity among translocated individuals.

**LAY SUMMARY:**

- The koloa maoli (Hawaiian duck, or koloa) is a federally-endangered, island endemic bird. Like other Hawaiian waterbirds, koloa are threatened by introduced predators and habitat loss, but also specifically by hybridization with feral mallards and localized avian botulism outbreaks. Currently, the island of Kaua‘i has the largest population of non-admixed koloa. We used capture-mark-recapture and genetic data of koloa at two wetland sites on Kaua‘i to examine their sociality and spatial genetic structure.
- Koloa formed preferential social associations, but they were not based on plumage traits, body mass or genetic relatedness.
- There was no spatial genetic structure for males and females within a wetland site. Our results suggested that 1) koloa have no preference for mallard-like plumage that might increase hybrid pairing, 2) localized (within-refuge) botulism outbreaks are unlikely to remove close relatives and unique genetic lineages, and 3) translocation of groups could maintain social associations without limiting genetic diversity.

## INTRODUCTION

Historically, island endemic birds had a greater rate of extinction than their mainland counterparts, and many extant island species are threatened with extinction (Fromm and Meiri 2021, Matthews *et al*. 2022). The loss of island endemic birds, which often have unique functional traits (Matthews *et al*. 2022), reduces both biodiversity and ecological richness. However, compared with their morphology and life-history, the sociality of island birds has been relatively underexplored (Jezierski *et al*. 2023). Island endemic birds experience unique ecological conditions that may impact social behaviors. For example, (1) non-native or depauperate predator communities may affect social grouping (Beauchamp 2004, 2010); (2) limited area and higher local densities (Blondel 2000) may increase opportunities for social interactions (Beck *et al*. 2024); (3) non-migratory behavior and small home range size provide the opportunity for longer-term affiliations (Reynolds *et al*. 2009); and 4) slower life-histories (i.e., high survival and low reproductive rates) may promote cooperative breeding (Covas 2012, Dibben-Young *et al*. 2021) and, consequently, social affiliations and/or genetic microgeographic structure (Sonsthagen *et al*. 2010, Bertrand *et al*. 2014). Hence, studies of island endemics are needed to identify potential unique social structures before they are lost.

Understanding the social behavior and emergent spatial genetic structure of endangered island endemics is also valuable for their recovery (Morris *et al*. 2021). For social species, translocations may disrupt bonds between conspecifics, thereby reducing survival and cohesion of translocated individuals, and ultimately decreasing translocation success (Goldenberg *et al*. 2019). Understanding links between social bonds and relatedness can inform the sourcing of individuals to be used in translocations and potentially preserve social relationships (Scribner *et al*. 2005, Taylor *et al*. 2021). Microgeographic spatial genetic structure reflects current and historical patterns in gene flow and may arise and be maintained in island birds due to local habitat heterogeneity (Cheek *et al*. 2022), landscape barriers to gene flow (Khimoun *et al*. 2017), reduced tendency to disperse by one or both sexes (Bertrand *et al*. 2014), or social including mate preferences (Langin *et al*. 2015). If kin are spatially clustered at small scales, localized pathogens or outbreaks are more likely to affect kin (e.g., chronic wasting disease in white-tailed deer (*Odocoileus virginianus*); Grear *et al*. 2010), which could result in correlated loss of genetic diversity.

Social network analysis has emerged as a useful and popular tool for analyzing animal societies and can provide critical insights for conservation and management (Snijders *et al*. 2017). As conservation actions are often resource-limited, repurposing existing data to examine additional questions is resource-efficient and reduces disturbance to animals. Many studies collect capture-mark-recapture (CMR) data, where individuals are uniquely marked and subject to recapture across multiple capture occasions. Although often used to evaluate population size and important vital rates, CMR data can also be used to infer social associations by assuming that marked individuals that are captured together are more likely to be associates of each other (Silk *et al*. 2021). For example, social networks of tigers (*Panthera tigris*) have been inferred from individuals sighted in the same location via camera traps and grouped within biologically meaningful time intervals (Carter *et al*. 2023). Similarly, genetic data collected for other objectives could also be used to infer spatial genetic structure.

The koloa maoli or Hawaiian duck (*Anas wyvilliana*; hereafter, koloa) is the only endemic duck species remaining in the main Hawaiian Islands (USFWS 2011). Although historically common, koloa experienced population declines and range contraction during the 20^th^ century (Banko 1987) and are listed as federally endangered (USFWS 2011). Modern hybridization with congeneric feral mallards (*Anas platyrhynchos*) led to their genetic extirpation from all islands except Kaua‘i and Ni‘ihau (USFWS 2011, Wells *et al*. 2019a), where their estimated population size is approximately 1,000-2,000 individuals, with a heavily male-biased adult sex ratio (Malachowski 2020, Paxton *et al*. 2021). On Kaua‘i, wetland complexes that are managed by the U.S. Fish and Wildlife Service (USFWS) on National Wildlife Refuges (NWRs) are heavily used by koloa (Malachowski 2020). A primary threat to this population includes highly localized outbreaks of Type C avian botulism, which are primarily proliferated via the carcass-maggot cycle wherein koloa become ill by feeding on fly larvae that have bioaccumulated the botulinum neurotoxin from toxic carcasses (Rocke and Bollinger 2007, Malachowski *et al*. 2022). Further, true (i.e., non-admixed) koloa on Kaua‘i exhibit variation in plumage, with some males exhibiting more mallard-like plumage traits, likely due to their ancestral hybrid origin (Lavretsky *et al*. 2015). Information on social associations and spatial distribution of kin could inform our understanding of plumage-based preferences for or against feral mallards and koloa-mallard hybrids, the potential for localized botulism outbreaks to remove entire lineages of relatives, and the planning of translocation efforts for koloa to other islands. Thus, our primary objectives were to (1) evaluate if koloa form non-random social associations and, if so, whether those social associations were influenced by more mallard-like plumage traits, body mass, or relatedness; and (2) determine whether koloa exhibit spatial genetic structure, possibly driven by female philopatry and male-biased dispersal.

## METHODS

### Study area

We conducted fieldwork at Hanalei and Hulē‘ia NWRs on the island of Kauaʻi, Hawaiʻi, USA. Hanalei NWR is a 371-ha refuge located on the north shore of Kaua‘i and is the most heavily used wetland site by koloa (USFWS 2011, Malachowski and Dugger 2018, Malachowski 2020). Site characteristics (e.g., land cover, vegetation, climate) at Hanalei NWR have been previously described (Malachowski and Dugger 2018, Malachowski *et al*. 2018); in brief, the refuge provided for 24 ha of management wetlands and 53 ha of farmed taro (*Colocasia esculenta*) ponds, or lo‘i, habitat (Malachowski and Dugger 2018). Hulē‘ia NWR is located along the Hulē‘ia river, ∼31 kilometers SW of Hanalei NWR, and covers 98 ha of river bottom habitat (USFWS 2011). Koloa breed throughout the year, with a peak between September-May (Malachowski *et al*. 2018, 2019). They face predation risk primarily from non-native species, such as cats (*Felis catus*), dogs (*Canis lupus familiaris*), rats (*Rattus* spp.), wild pigs (*Sus scrofa*), American Barn Owls (*Tyto furcata*), western cattle egrets (*Bubulcus ibis*), and American bullfrogs (*Lithobates catesbeianus*) (Malachowski *et al*. 2019, 2022, Webber 2022).

### Capture and sampling

We captured, banded, and processed birds at Hanalei NWR during December 2010–November 2015 (see Supplementary Material Table S1 for sampling schedule) and Hulē‘ia NWR during July–August 2011 using methods previously described (Malachowski 2020; Malachowski *et al*. 2022). In short, we captured birds using customized baited swim-in traps (BST) placed in managed wetland units. We banded each bird with a U.S. Geological Survey (USGS) metal leg band, determined sex and age (duckling, juvenile, or first-year/adult [hereafter, adult]), and collected morphometric data (e.g., body mass). We assigned males a combined plumage score based on four binary plumage traits that corresponded to 1) percentage of green on head, 2) chest color and pattern, 3) belly and flank pattern, and 4) back pattern (Supplementary Material Table S2); plumage scores ranged from 0 (no mallard-like traits) to 4 (all mallard-like traits). Although most koloa on Kaua‘i are not admixed (Wells *et al*. 2019a; Fukunaga *et al*. 2026), koloa express variation in plumage, including mallard-like traits, presumably because of their ancestral hybrid origin from Laysan ducks (*Anas laysanensis*) and ancestral wild mallards (Lavretsky *et al*. 2015). Last, we collected blood samples from a subset of koloa during March–August 2011 and obtained mitochondrial haplotypes and single nucleotide polymorphism genotypes at 3,308 loci (see Wells *et al*. 2019a). For all recaptured birds, we recorded USGS band numbers and, if recaptured ≳ 1 month from its previous capture, we re-sampled morphometric data and plumage scores.

For subsequent analyses, we considered individuals captured in the same trap during the same trap-night (i.e., dusk to dawn) to be part of the same ‘group’ and associates of each other. We only included data from trapping events where the identities of all captured individuals were known. Because we had too few ducklings and juveniles to consider as separate age classes, we only used associations among adults for all analyses. Given sample size constraints at Hulē‘ia, we only used data for birds captured at Hanalei for all analyses, unless otherwise noted.

### Data analysis

#### Preferential associations among koloa

We tested for preferential (or non-random) associations among male-male (MM), male-female (MF), and female-female (FF) koloa dyads (i.e., pairs of individuals) using a sampling period of 30 days, as mortality is low within this timespan and trapping events typically lasted <30 days after 2011. To do this we compared observed associations with randomized associations using several types of permutations (i.e. randomizing groups or associations while keeping group sizes and sampling structure constant); we used SOCPROG v 2.9 (Whitehead 2009) for all analyses. We first used the ‘*permute groups within samples*’ permutation method, where the elements of the groups-by-individual incidence matrix (1 if an individual was present in the group, 0 if not) in each 30-day sampling period is permuted, while keeping the row and column totals constant. This method accounts for individuals missing across sampling periods and the number of groups with which individuals associated (which is the number of times individuals were captured) within sampling periods but does not control for differences in individual gregariousness. If any of the dyad datasets (MM dyads, MF dyads, or FF dyads) showed significant variation in individual gregariousness, we controlled for gregariousness by repeating the analysis with the ‘*permute associations within samples*’ method, where the association matrix between individuals is permuted in each sampling period, keeping row and column totals constant.

We used the half-weight index as the measure of dyadic associations because it limits bias when all individuals within each group have not been captured and identified (Whitehead 2008). This approach was warranted because ducks in a trap were considered a sample of all ducks using the associated wetland impoundment on a given night due to factors such as individual heterogeneity in capture probability (Malachowski 2020) and limitations of trap size. For example, koloa occasionally gather in large groups (e.g., >100 birds; Weller 1980, USFWS 2011, Malachowski 2013) which exceeds the physical capacity of a trap. Because male and female koloa have different capture probabilities (Malachowski 2020), we also repeated MF dyad analysis with the social affinity index, which is less biased when there are differences in capture probabilities between categories (Whitehead 2008). We performed permutations of the three datasets (MM, MF, FF), starting with 10,000 permutations and increasing the number of permutations in increments of 10,000 until the *P-*values obtained were similar (Whitehead 2009); we report results based on 20,000 permutations. A significantly higher coefficient of variation (CV) of observed association indices compared with those generated from random datasets (i.e., observed CV greater than 95% of the CV values from random datasets) suggests the presence of preferential associations that persist across sampling periods (Whitehead 2009).

#### Calculating association indices and constructing networks

We calculated half-weight association indices (AI) between individuals captured ≥ 3 times in the entire dataset and constructed networks of associations using Gephi 0.10 (Bastian *et al*. 2009); these AI values were used in the Mantel tests and node-label permutations described below. AI varies from 0 (i.e., pairs never captured together) to 1 (i.e., pairs always captured together). Because of mortality and emigration (Malachowski 2020; Malachowski *et al*. 2022), combining all individuals captured over five years might include dyads who were never present at the same time on the refuge. Therefore, we also constructed the network and repeated all Mantel tests (Mantel 1967) and node-label permutations using ducks captured ≥ 3 times during a two-year period (2011-2012) with the greatest number of annual captures (Supplementary Material Table S1).

#### Relationship between associations and male plumage traits and koloa body mass

Koloa plumage scores appear to vary seasonally and among individuals, with some males exhibiting higher scores (i.e., more mallard-like plumage) during November–April (Wells *et al*., unpublished data). To best capture these differences among males we used scores and captures (used to calculate AI values) from November to April for plumage analyses. If a male had multiple plumage score measurements in that dataset, we used the average score, but results were similar when instead using the maximum score.

To examine whether males showed assortative associations based on plumage, we evaluated the relationship between AI and the absolute difference in their plumage scores using Mantel tests with Spearman’s rank correlation and 1,000 permutations. A negative correlation would indicate males associated more with other males with similar plumage scores. All Mantel tests were performed using the mantel function in the *ecodist* package (Goslee and Urban 2007). We also tested whether males with higher plumage scores were preferred associates of females and other males by assessing their weighted degree, calculated as the sum of edge weights (in this case, AIs) linking a given male to other individuals. The statistical significance of the Pearson’s correlation coefficient between the average plumage score of a male and his weighted degree was determined using a permutation test with 1,000 node-label permutations, where we randomly shuffled average plumage scores across the males in the network to create permuted datasets, and compared the observed correlation to this null distribution (Farine 2017, Weiss *et al*. 2021). As we had no *a priori* expectation for the direction of the relationship between plumage score and weighted degree, we used a two-tailed test, where we considered observed correlations to be significant if they were ≤2.5^th^ or ≥97.5^th^ percentiles of the corresponding permuted correlation coefficients.

We similarly used Mantel tests to evaluate the relationship between AI of same-sex dyads and the absolute difference in their average mass. A negative correlation would indicate koloa associate more with conspecifics with similar mass; this might occur if similarly-sized koloa formed groups due to similar activity budgets, which could help group cohesion (Ruckstuhl and Neuhaus 2002), or if some individuals who prefer to feed with larger individuals (Duffy *et al*. 2009) benefit from that association and are heavier themselves. To examine whether heavier individuals were better connected in the overall network, we used permutation tests with 1,000 node-label permutations, as previously described (average body mass values were randomly shuffled between individuals of the same sex in the network), to compare observed correlations between average mass and weighted degree to the null distributions for MM, MF, FF, and FM dyads. If heavier individuals were preferred as associates by others (e.g., successful foragers are heavier and preferred by all others as associates; Duffy *et al*. 2009), we would expect a significant positive correlation between mass and the network metrics. If they were avoided (e.g., lighter individuals are subordinates and minimize risk of receiving aggression by avoiding heavier individuals; Madden *et al*. 2011), we would expect a negative correlation.

#### Relatedness, microspatial genetic structure and female-biased philopatry in Hanalei

We used the DyadML estimator (see Supplementary Material Table S3 for details) in the Coancestry program v.1.0.1.10 (Wang 2011) to calculate relatedness for the 93 koloa from Hanalei NWR and 15 from Hulē‘ia NWR with complete genetic data (1502 variable loci with no missing data and one of 10 mitochondrial haplotypes; Wells *et al*. 2019a). We examined the correlation between dyad relatedness and AI using Mantel tests with Spearman’s rank correlation and 1000 permutations for MM and FF dyads separately. A positive correlation between relatedness and AI would suggest that individuals are more likely to associate with relatives than with unrelated individuals.

To test if haplotype similarity influenced association patterns, we calculated the expected proportion of MM, MF, and FF dyads with different mitochondrial haplotype frequencies. Since many haplotypes were rare, we simplified the data by grouping them into two categories: the most common haplotype and an ‘other’ category, which pooled all other haplotypes. This resulted in three possible dyad types: 1) both individuals share the common haplotype; 2) one individual has the common haplotype, and the other has an ‘other’ haplotype; or 3) both individuals have ‘other’ haplotypes. Then, we used Chi-squared tests to compare the observed number of associated haplotyped dyads in each of the three haplotype categories with the frequencies that would be expected by chance. If individuals with the same haplotype preferentially associated with each other, we expected the observed proportions of same- haplotype dyads among associating pairs to be greater than expected frequencies.

We calculated pairwise distances between capture locations of all genotyped ducks captured within 3-month periods, when there were sufficient numbers of captures of genotyped koloa. If the trap location for a given capture record was uncertain between two locations (*n* = 20), we used the midpoint. For ducks captured multiple times during a 3-mo period, we used the average of its capture location coordinates to calculate its distance to other ducks. We examined the effect of distance and the interaction of distance and dyad type (MM, MF, FF) on pairwise relatedness (*r*), with the time period as a random effect, using a zero-inflated generalized linear mixed model with a log-normal distribution and a log link function. If there was spatial genetic structure within Hanalei NWR, we expected dyads captured farther apart to be less related than those captured at shorter distances. If there was male-biased dispersal, we expected relatedness to decrease with distance for FF dyads, but not necessarily for MM dyads. Further, if spatial sub- structuring was important for inbreeding avoidance, we expected related MF dyads to be captured farther apart than unrelated ones. Similarly, we examined the effect of distance and its interaction with dyad type on whether a dyad had the same (0) or different (1) haplotypes, using a generalized linear mixed model with a binomial distribution and a logit link and treating time period as a random effect. We implemented models and validated assumptions using packages glmmTMB (Brooks *et al*. 2017) and DHARMa (Hartig 2021), respectively, in Program R (version 4.2.3, R Foundation for Statistical Computing, Vienna, Austria).

To evaluate possible female-biased philopatry at Hanalei NWR, we tested whether FF dyads in Hanalei had higher average relatedness than MM and MF dyads using the ‘Test Group Difference’ option in Coancestry Version 1.0.1.10 (Wang 2011) with 1,000 bootstraps to determine significance. The dyad type categories being compared are shuffled together to create randomized datasets with the same number of dyads as the original datasets, and the observed difference in average relatedness are compared with differences in random datasets. If the observed difference was greater than the 99% quantile of the distribution of differences in the random datasets, it is significant. Similarly, we examined sex differences in haplotypes within Hanalei by comparing the haplotype counts of adult females with haplotype counts of adult males using a G-test of independence with Williams’ correction (Sokal and Rohlf 1981). Given B-type haplotypes are typically found in koloa (Fowler *et al*. 2009), we excluded individuals with A-type haplotypes (i.e., those found in mallards) and pooled all rare haplotypes that occurred in less than four individuals. If females were more philopatric than males within Hanalei, we expected 1) FF dyads in Hanalei would be more related than MM and MF dyads, and 2) haplotypes of males would be different than females.

#### Regional-scale relatedness and genetic structure

To investigate genetic structure at a larger spatial scale, we first examined if any closely related adults (*r* ≥ 0.125) were captured at different refuges (i.e., between-refuge MM, MF, and FF dyads). We also compared the relative frequencies of haplotypes of all individuals between refuges using a G-test of independence with Williams’ correction (Sokal and Rohlf 1981). We included individuals captured by all methods (not just the BST method) and again excluded A-type haplotypes and pooled all rare haplotypes (occurred in less than four individuals). If females are more philopatric than males at the refuge scale, we expect a lower proportion of closely related between-refuge FF dyads than MM and MF dyads. Alternatively, if there is little or no movement of koloa of both sexes between the refuges, we expect to find no closely related between-refuge dyads and differences in haplotype frequencies between the refuges.

## RESULTS

We recorded 267 trapping events (i.e., groups) that met our sampling criteria between December 2010 and November 2015 at Hanalei NWR. This included 902 individuals (*n*_male_ = 730; *n*_female_ = 172) that were captured 1,659 times. Adult group size (i.e., number of adults captured together) varied from 1 to 27 (Avg. ± SD: 6.2 ± 5.4; mode = 1; Figure 1a). The average number of captures per individual was low (Females- Maximum: 6, Avg. ± SD: 1.5 ± 0.9; Males- Maximum: 12, Avg. ± SD: 1.9 ± 1.5). Most male and female koloa moved between wetland impoundments (or units) between captures; 73% of males and 71% of females captured at least three times were captured in more than one unit (Figure 1b), and of those koloa captured more than 30 days apart (n = 109), 93% were captured in more than one unit.

**Figure 1.**
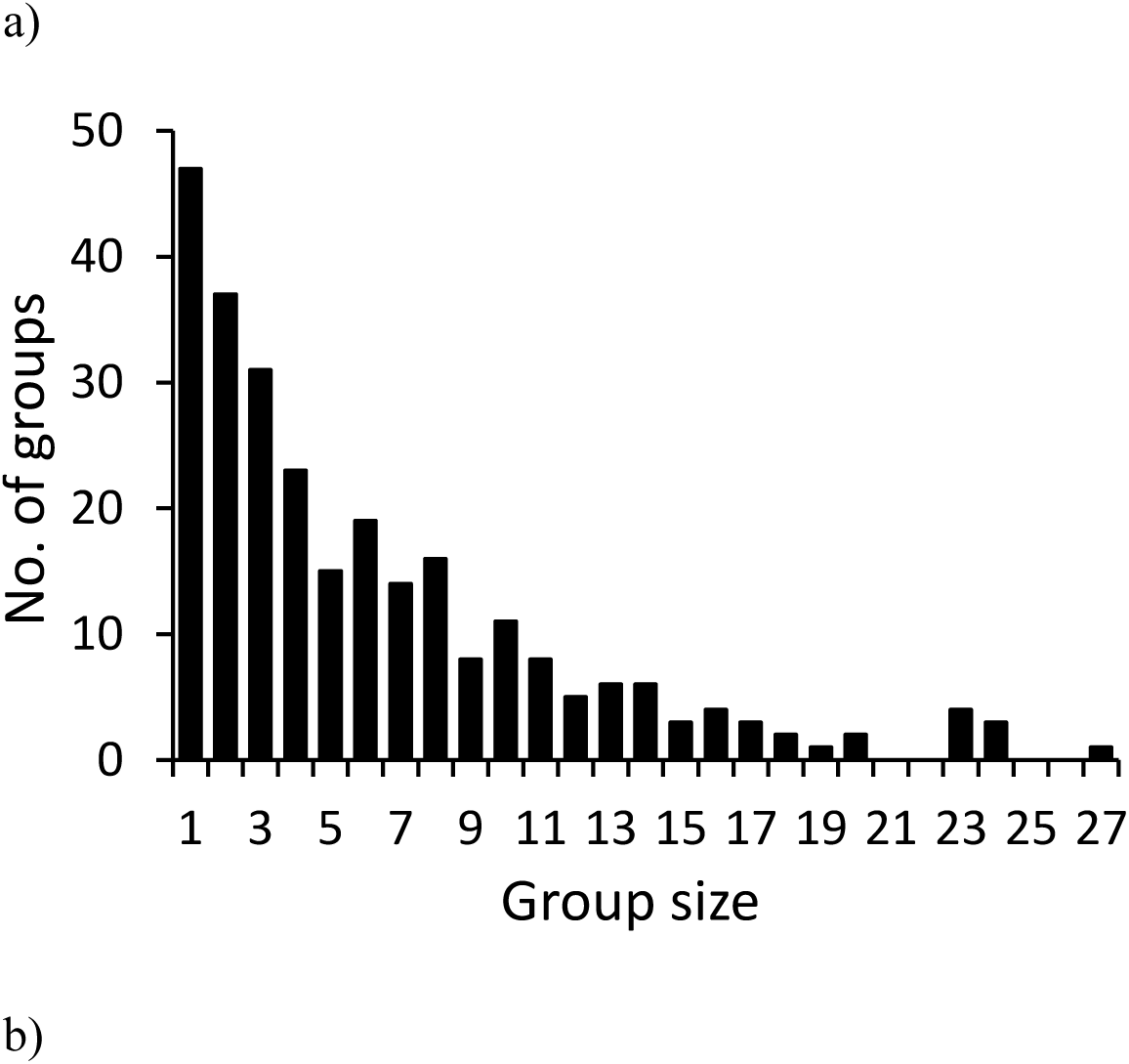
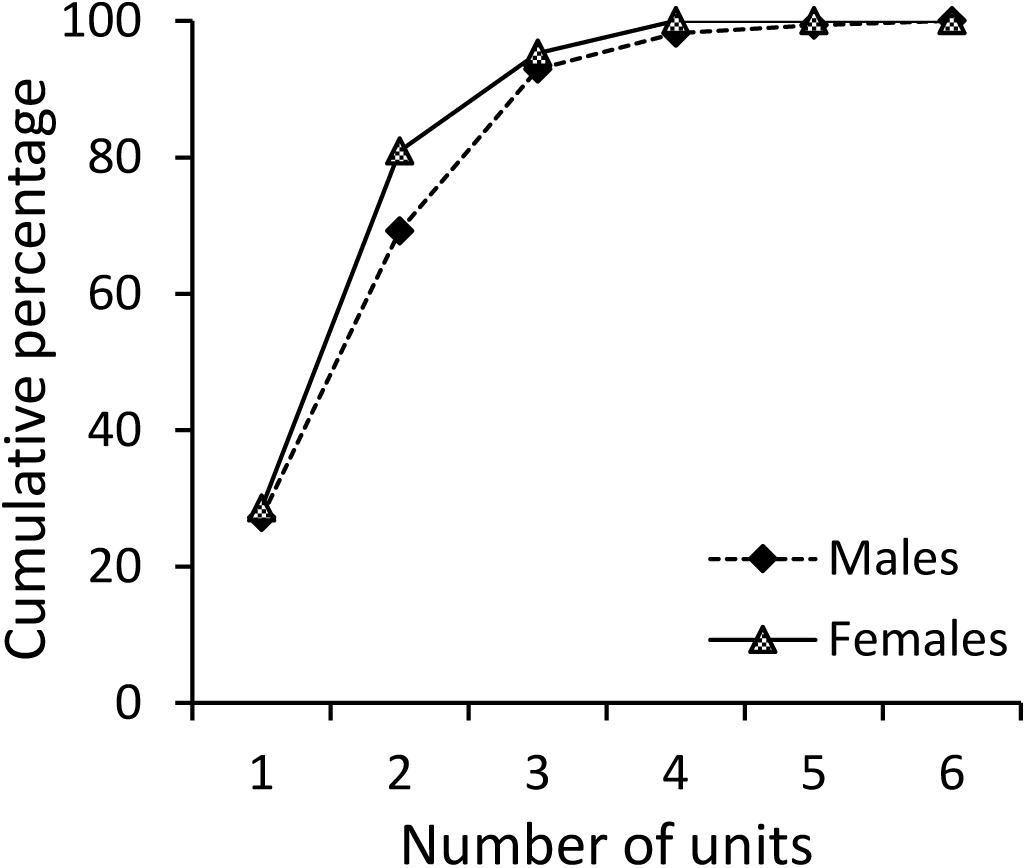
a) Histogram of adult koloa maoli group sizes (i.e., number of adults captured together in a baited swim-in trap; *n* = 267 groups) during December 2010–November 2015 at Hanalei National Wildlife Refuge, Kaua‘i, Hawai‘i USA. b) Cumulative percentage of males (*N* = 169) and females (*N*= 21) of adult koloa maoli captured at Hanalei National Wildlife Refuge, Kaua‘i, Hawai‘i, USA, during December 2010–November 2015, and the number of units (wetland impoundments within the refuge) in which they were captured.

**Figure 2.**
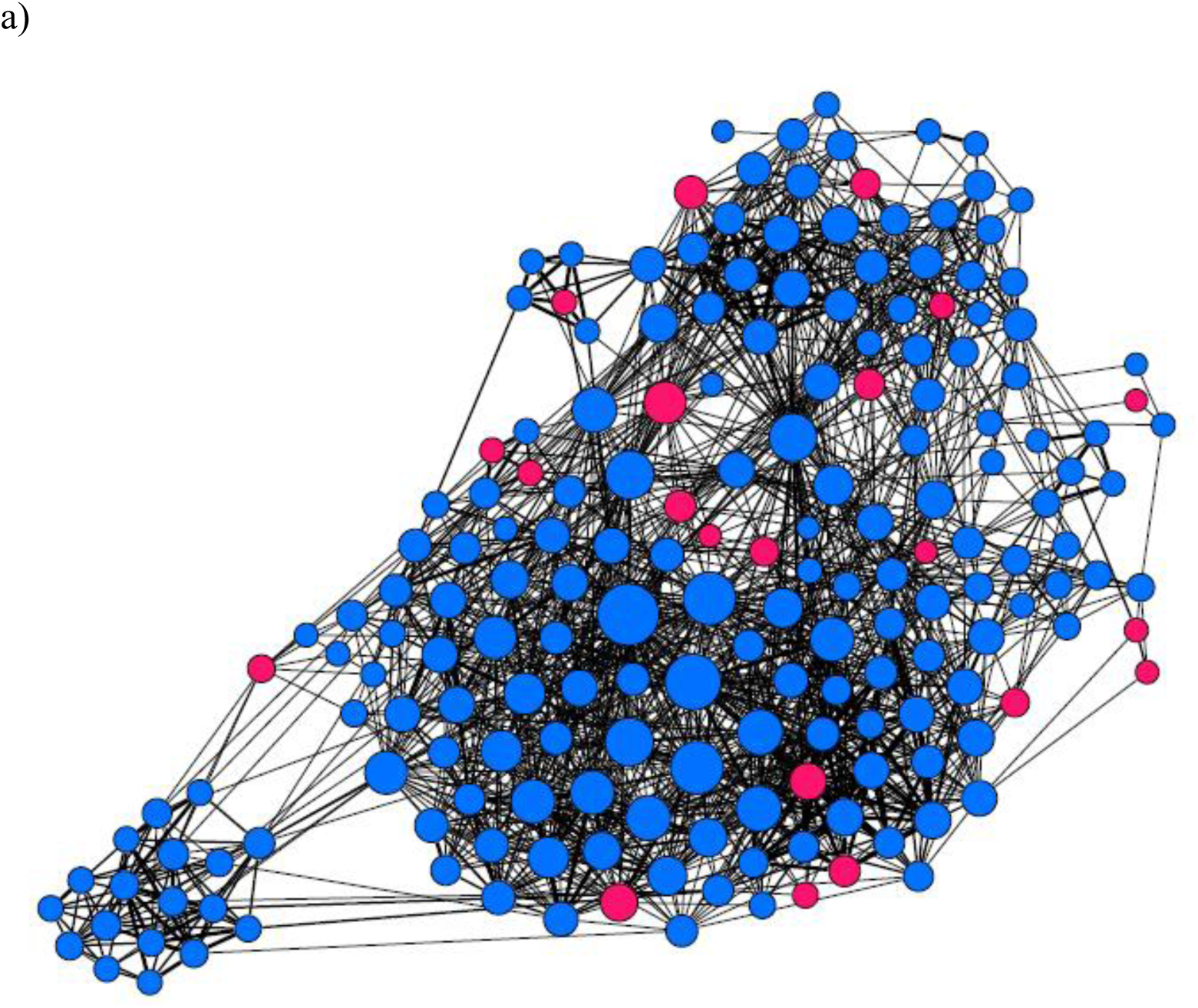
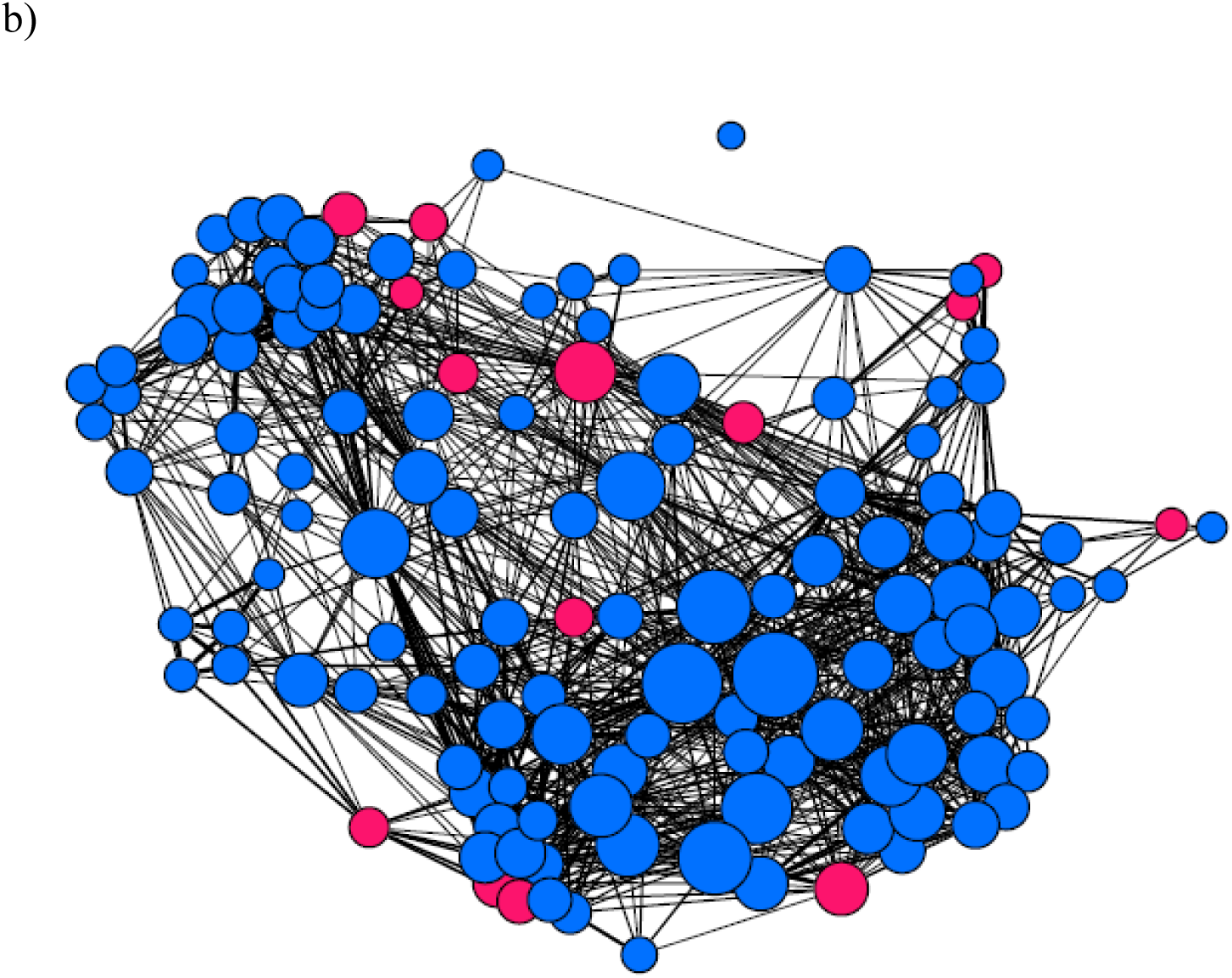
Networks of associations among individual male (blue nodes) and female (pink nodes) adult koloa maoli captured at Hanalei National Wildlife Refuge, Kaua‘i, Hawai‘i, USA, during a) December 2010–November 2015, and b) January 2011–December 2012. The nodes are sized by their degree (i.e., number of connections or associates), and edges (i.e., lines connecting pairs of nodes) are weighted by their half-weight association index, where thicker edges represent stronger associations.

### Preferential associations among koloa

Using the half-weight index, males exhibited preferential associations across sampling periods with females and other males (Table 1; results for MF dyads remained consistent with the social affinity index, see Supplementary Material Table S4). By contrast, females did not show preferential associations with other females (Table 1). Only MM dyads showed significant variation in individual gregariousness (SD_observed_ = 5.7188; S̅̅D̅̅_permuted_ = 5.5050; *P* = 0.0060), but their preferential associations remained significant after controlling for differences in individual gregariousness (Supplementary Material Table S4).

**Table 1.** Results of tests for preferential associations between male-male (MM), male-female (MF), and female-female (FF) dyads for adult koloa maoli captured together at Hanalei National Wildlife Refuge, Kaua‘i, Hawai‘i, USA, during December 2010–November 2015. We compared the coefficient of variation (CV) of half-weight association indices of the observed dataset with values from 20,000 permuted datasets and calculated *P* as 1 minus the proportion of permutations where CV_observed_ > CV_permuted_. Permuted datasets were created by permuting the groups-by-individual incidence matrix in each 30-day sampling period, while keeping the row and column totals constant.

| Dyad | $CV_{\text{observed}}$ | Mean of $CV_{\text{permuted}}$ | $P$ |
| --- | --- | --- | --- |
| MM | 7.1258 | 7.0011 | **<0.0001** |
| MF | 9.0189 | 8.9179 | **<0.0001** |
| FF | 10.2386 | 10.1686 | 0.1096 |

### Relationship between associations and male plumage traits and koloa body mass

We found no evidence for assortative associations among males based on plumage score either in the 5 year or shorter (2011–2012) dataset (Table 2). We also found no evidence that males with higher plumage scores were preferred associates of females or other males based on weighted degrees during either time period (Table 3). Similarly, we found no significant relationships between AI of same-sex dyads and absolute difference in body mass (Table 2), nor between individuals’ body mass and their weighted degree within and across sexes during either time period (Table 3).

**Table 2.**
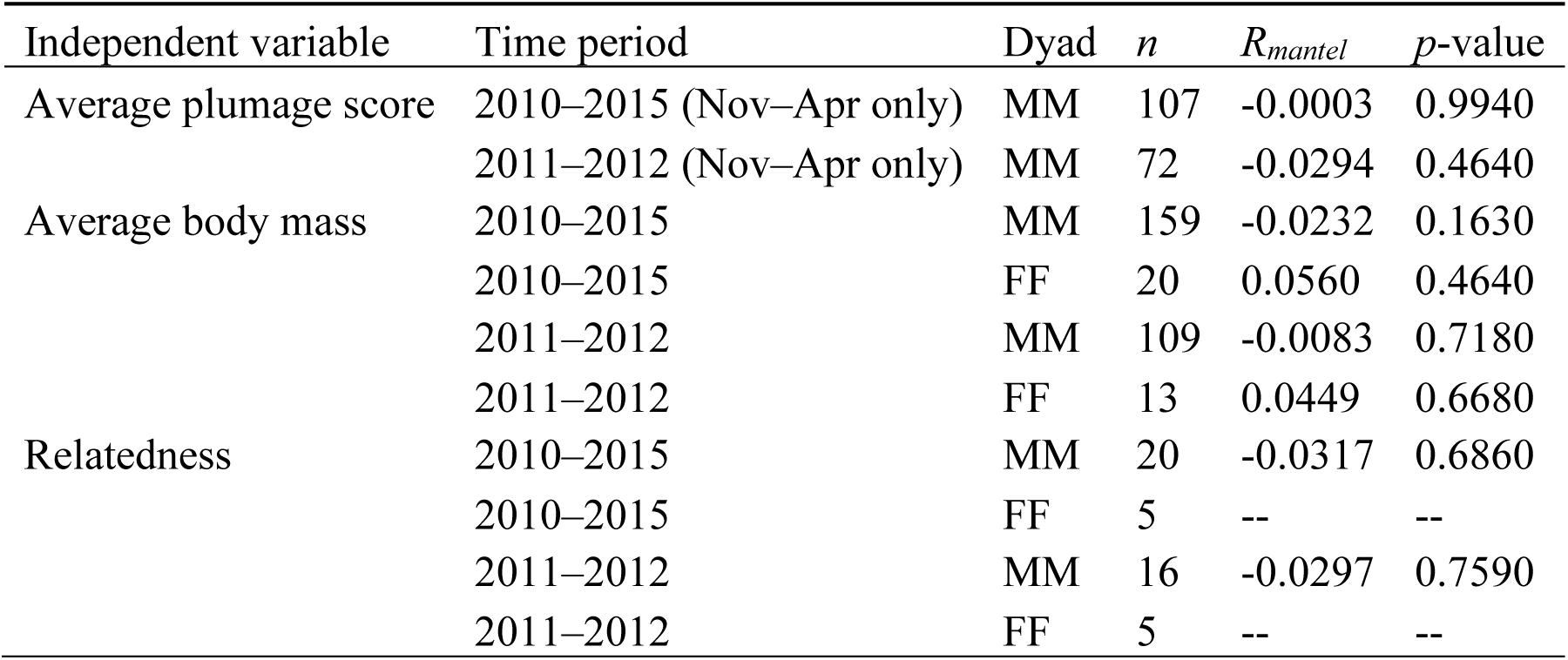
Relationships between half-weight association index and 1) absolute difference in plumage scores, 2) absolute difference in average body mass, and 3) genetic relatedness for male-male (MM) and female-female (FF) dyads for adult koloa maoli captured together at Hanalei National Wildlife Refuge, Kaua‘i, Hawai‘i, USA, during December 2010–November 2015. We used Mantel tests with Spearman’s rank correlation (*R_mantel_*) and 1,000 permutations, and determined significance using a two-tailed test with a significance level of 0.05. Raw data for all comparisons are presented in Supplementary Material Figure S2.

**Table 3.**
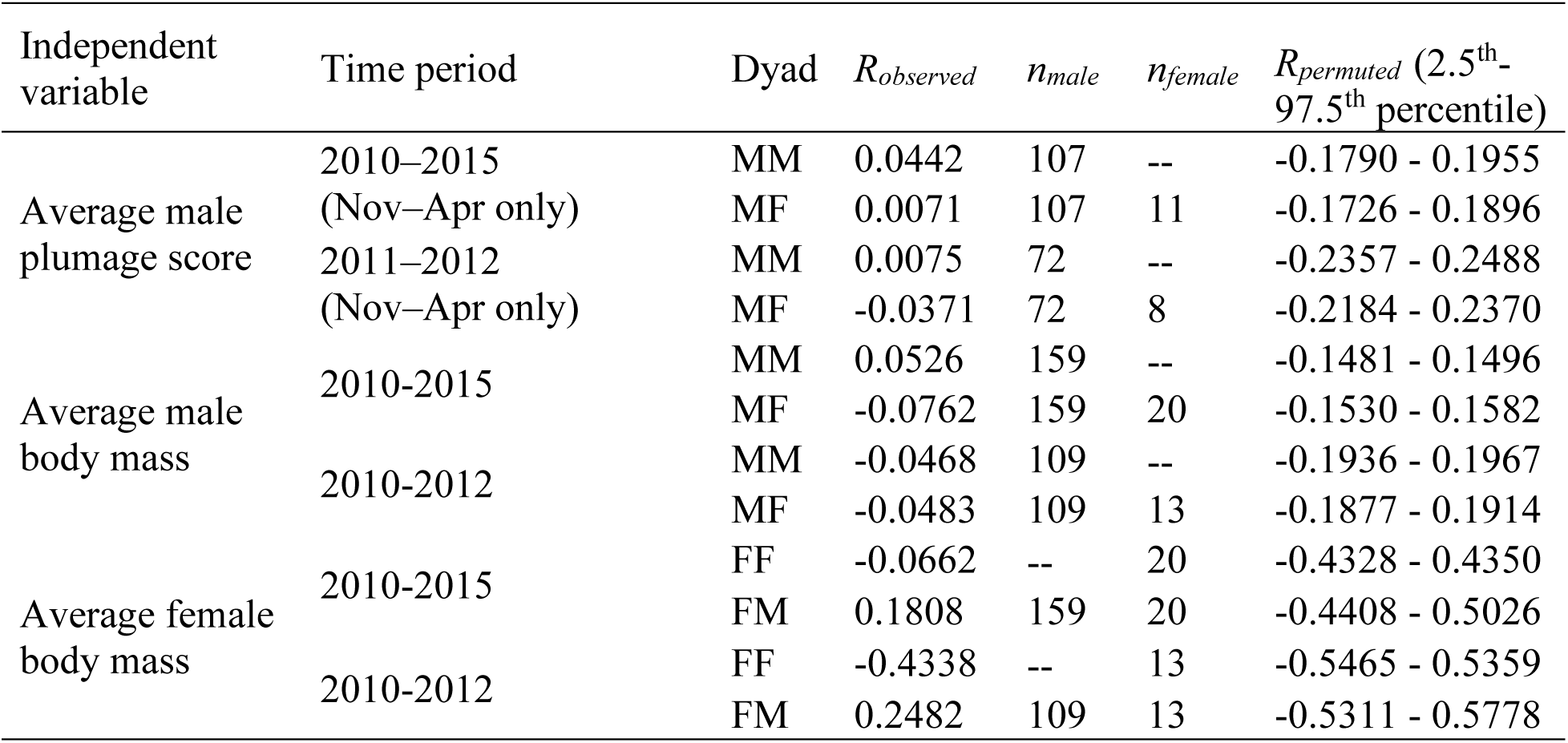
Relationships between the weighted degree (sum of AI) of 1) males with other males and females and his average plumage scores, 2) males with other males and females and his average body mass, and 3) females with other females and males and her average body mass, for adult koloa maoli captured together at Hanalei National Wildlife Refuge, Kaua‘i, Hawai‘i, USA, during December 2010–November 2015. We compared observed Pearson’s correlation coefficient with those in 1000 permuted networks. Networks were permuted by node label permutations, where plumage scores and mass values were shuffled randomly between individuals. The observed correlation was significant if it was ≤2.5^th^ or ≥97.5^th^ percentiles of the corresponding permuted coefficients. Raw data for all comparisons in Supplementary Material Figure S3 and Supplementary Material Figure S4.

### Relatedness, microspatial genetic structure and female-biased philopatry in Hanalei

Average dyad relatedness among the 108 genetically sampled koloa (*n*_Hanalei_ = 93; *n*_Hulē‘ia_ = 15) was low, but there were a few closely related dyads (*n* = 5,778 possible dyads; 𝑟̅ ± SD: 0.0107 ± 0.0334, maximum *r* = 0.5726). Of the 10 different haplotype subgroups found on Hanalei, 2 were of the A-type, and 8 were of the B-type. Genotyped dyads captured in the same trapping event (associates) mostly had low relatedness, with only one MM dyad with *r* > 0.50 (Figure 3). We found no evidence that males preferentially associated with more related males (Table 2). We could not statistically compare relatedness or haplotype with AI among females because of low sample size (*n* = 8; Table 2). Most (73.0% of 63 males, and 92.3% of 13 females) haplotyped koloa at Hanalei had Haplotype 1 (type B haplotype that corresponds to Haplotype J in Fowler *et al*. 2009). Only two individuals had an A- type haplotype. Haplotypes of associating dyads were not different from the expected values based on overall haplotype frequencies for MM (*n* = 148 associating dyads, χ^2^ = 5.2389, *P* = 0.0728), MF (*n* = 46, χ^2^ = 4.6775, *P* = 0.0964) or FF (*n* = 4, χ^2^ = 0.6944, *P* = 0.7066) dyads

**Figure 3.**
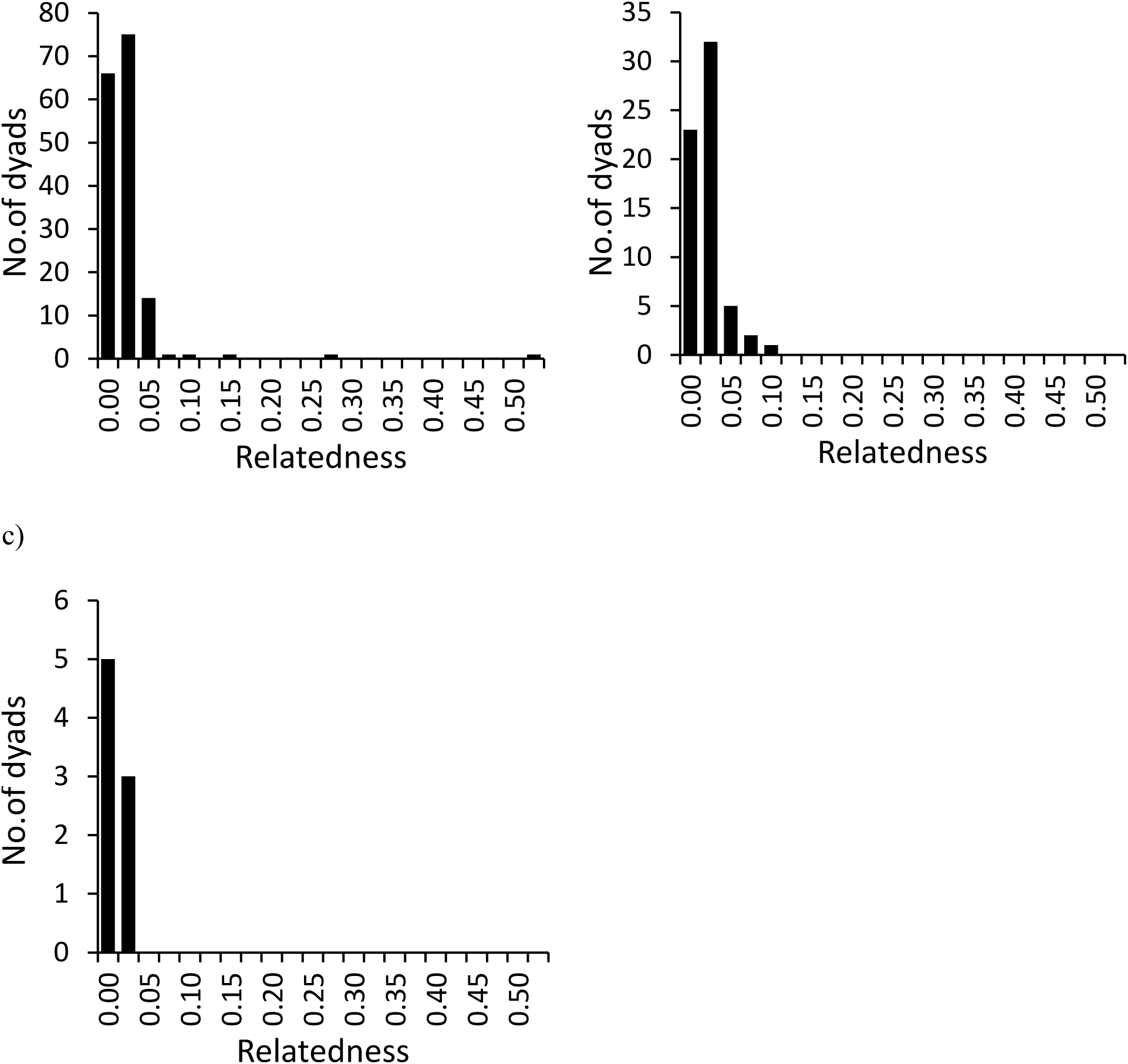
Histogram of relatedness (*r*) for a) male-male (*n* = 160), b) male-female (*n* = 63), and c) female-female (*n* = 8) dyads for adult koloa maoli captured together at Hanalei National Wildlife Refuge, Kaua‘i, Hawai‘i, USA, during December 2010–November 2015.

**Figure 4.**
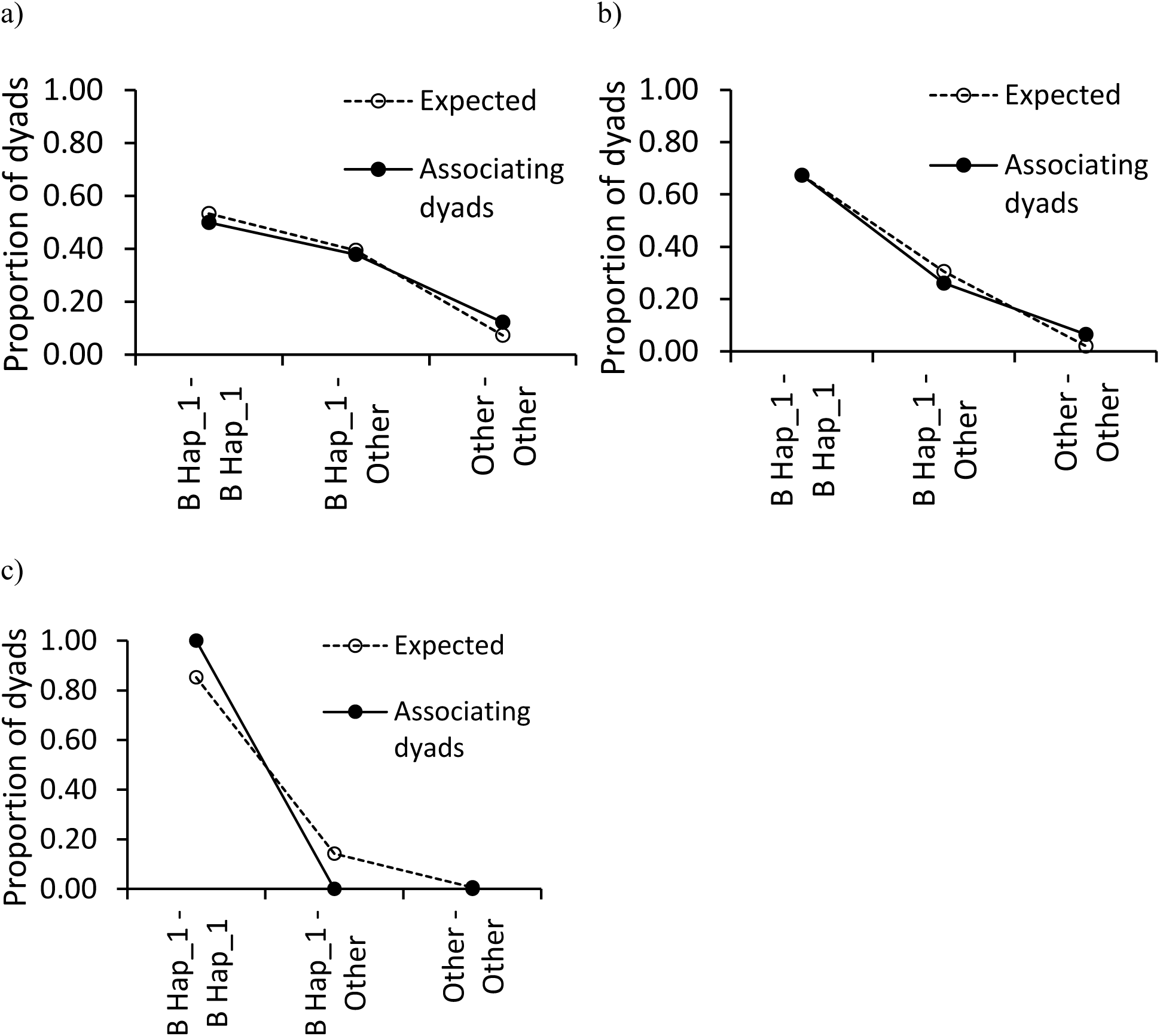
Expected (based on overall haplotype frequencies of male and female koloa on Hanalei) and observed proportion of associating dyads (dyads of adult koloa maoli that were captured together) with different haplotype combinations in a) male-male dyads (*N* = 148), b) male-female dyads (*N* = 46), c) female-female dyads (*N* = 4) captured at Hanalei National Wildlife Refuge, Kaua‘i, Hawai‘i, USA, during December 2010–November 2015. Other haplotypes include Haplotype subgroups 2, 3, 4, 5, 6, 7, 8, 19, and 20. Haplotype subgroups 4 and 7 are A-type (more common in mallards) subgroups, and only two koloa in this dataset had these haplotypes.

Based on 1,591 genotyped koloa dyads, pairwise distances between average capture locations within five different 3-month periods averaged 296 ± 302 (SD) m, and relatedness averaged 0.0122 ± 0.0404 (SD). We found no significant effects of distance or distance × dyad type interaction terms on relatedness (all *p*-values > 0.05; Table 4, Figure 5; relatedness values binned by distance in Supplementary Material Figure S5). Pairwise distances between average capture locations within 3-month periods of dyads with haplotype data averaged 257 ± 271 (SD) m for dyads with the same haplotype (*n* = 728 dyads) and 331 ± 332 (SD) m for dyads with different haplotypes (*n* = 630 dyads). We found no evidence that distance or its interaction with dyad type affected whether the dyad had the same or different haplotypes (all *p*-values > 0.05; Table 4, Figure 6).

**Figure 5.**
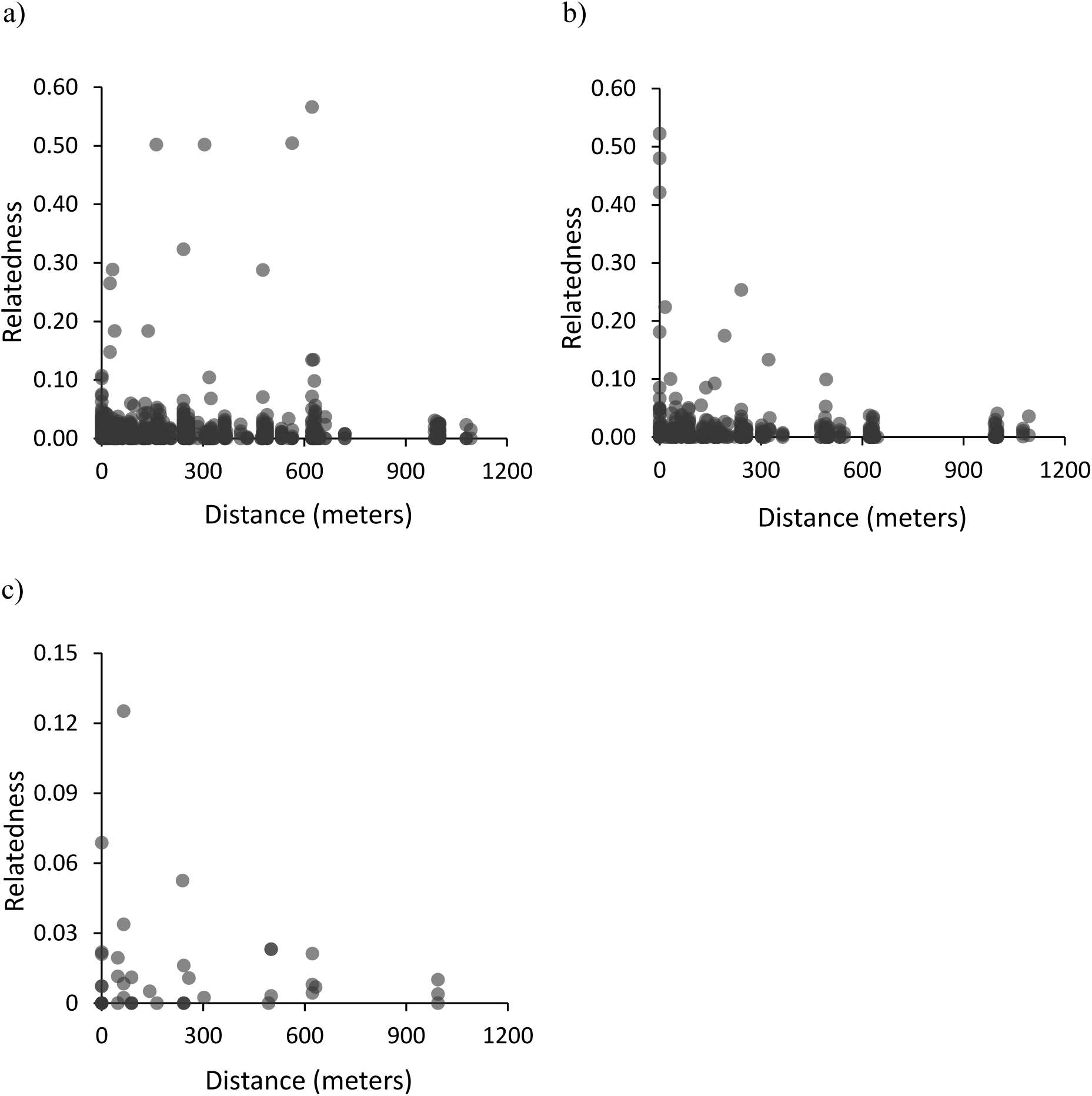
Scatter plots of distance between dyads of adult koloa maoli which were captured at Hanalei National Wildlife Refuge, Kaua‘i, Hawai‘i, USA, within the same 3-month period during December 2010–November 2015, and the relatedness between them for a) male-male dyads (*N* = 1070), b) male-female dyads (*N* = 478), c) female-female dyads (*N* = 43).

**Figure 6.**
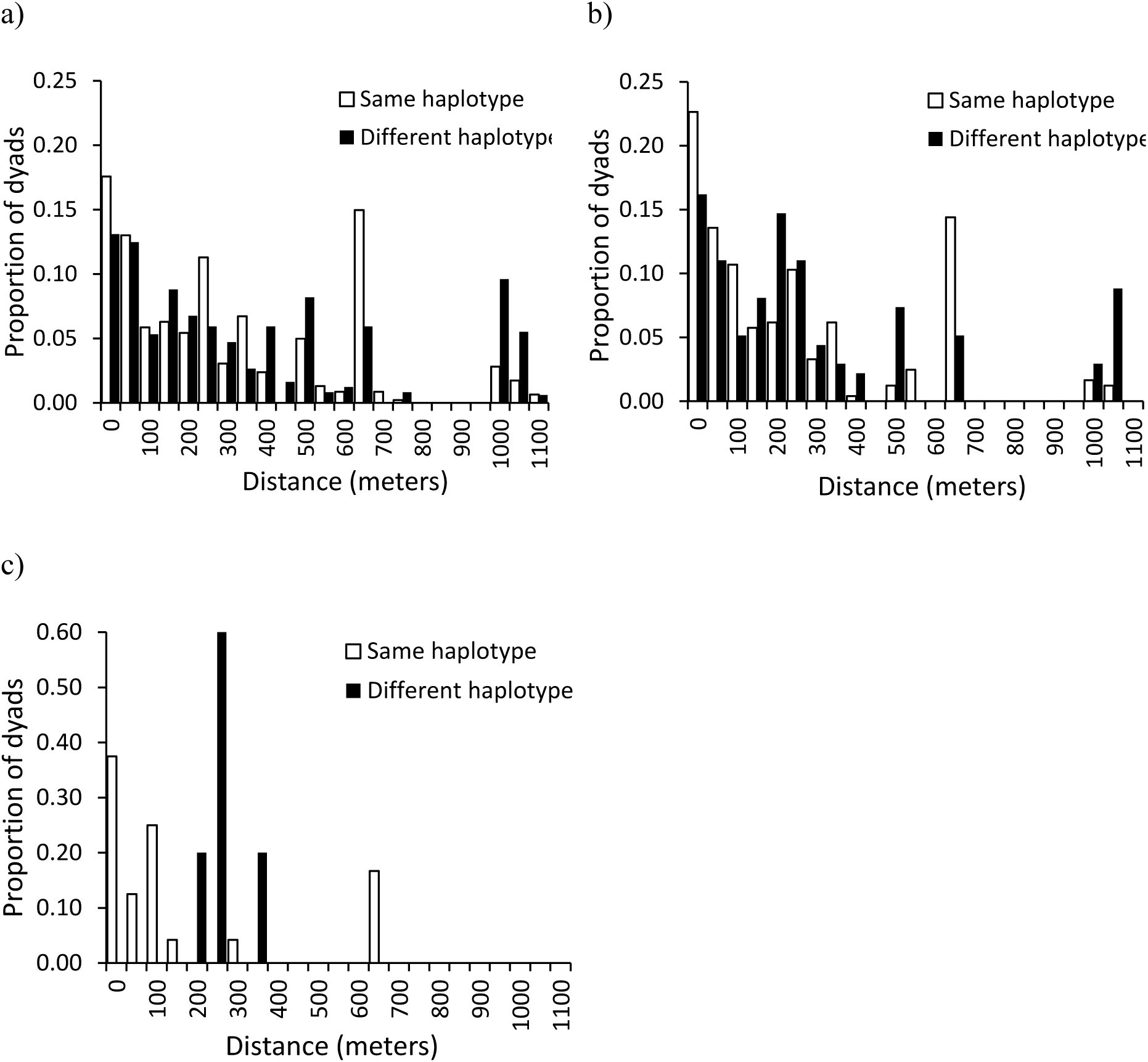
Distances between dyads of adult koloa maoli which were captured at Hanalei National Wildlife Refuge, Kaua‘i, Hawai‘i, USA, within the same 3-month period during December 2010–November 2015, with the same and different haplotypes for a) male-male dyads (*N_same_* = 461, *N_different_* = 489), b) male-female dyads (*N_same_* = 243, *N_different_* = 136), c) female-female dyads (*N_same_* = 24, *N_different_* = 5).

**Table 4.** Results of the generalized linear mixed models examining how distance between average capture locations within 3-month periods and dyad type (male-male, MM; male-female, MF; and female-female, FF, used as the reference dyad) influence relatedness (r∼Distance*Dyad_Type -Dyad_Type +(1|Period)) and haplotype similarity (Haplotype∼Distance*Dyad_Type -Dyad_Type +(1|Period)) in dyads of adult koloa maoli which were captured at Hanalei National Wildlife Refuge, Kaua‘i, Hawai‘i, USA, within the same 3- month period during December 2010–November 2015.

| Dependent variable | Fixed effects | Coefficient ± SE | z-value | P |
| --- | --- | --- | --- | --- |
| Relatedness | Intercept | -3.9168 ± 0.0569 | -68.8200 | **<0.0001** |
| | Distance | 0.0002 ± 0.0002 | 0.9700 | 0.3300 |
| | Distance : MF dyads | -0.0003 ± 0.0002 | -1.4500 | 0.1480 |
| | Distance : MM dyads | -0.0003 ± 0.0002 | -1.4000 | 0.1610 |
| | Random effects | Variance (±sd) | | |
| | Period | 1.331e-10 ± 1.154e-5 | | |
| Haplotype | Intercept | -0.3959 ± 0.1029 | -3.8490 | **<0.0001** |
| | Distance | -0.0017 ± 0.0018 | -0.9400 | 0.3472 |
| | Distance : MF dyads | 0.0018 ± 0.0018 | 1.0000 | 0.3172 |
| | Distance : MM dyads | 0.0028 ± 0.0018 | 1.5670 | 0.1171 |
| | Random effects | Variance (±sd) | | |
| | Period | 0.0109 ± 0.1045 | | |

### Among the subset of genotyped ducks that were captured as adults on Hanalei (*n*_male_ = 65; *n*_female_

= 14), average relatedness of FF dyads was not greater than MM dyads (observed difference (FF- MM): -0.0022; 99% quantile of differences in randomized datasets: 0.0102) or MF dyads (observed difference (FF-MF): -0.0025; 99% quantile of differences in randomized datasets: 0.0116). We also found no difference in haplotype frequencies between adult males and adult females (*G_adj_* = 4.3062, *df* = 3, *P* = 0.2302; Figure 7a).

**Figure 7.**
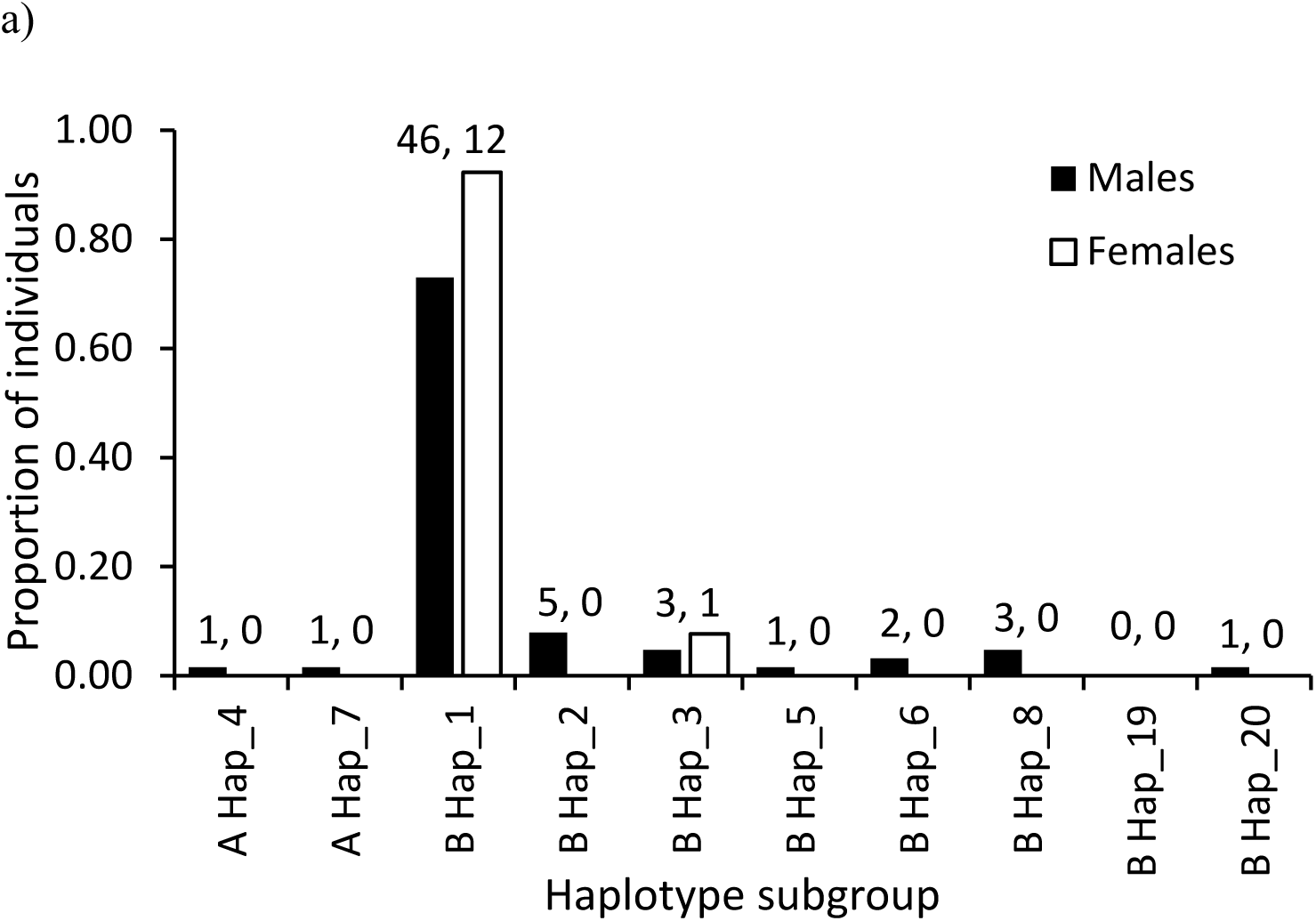
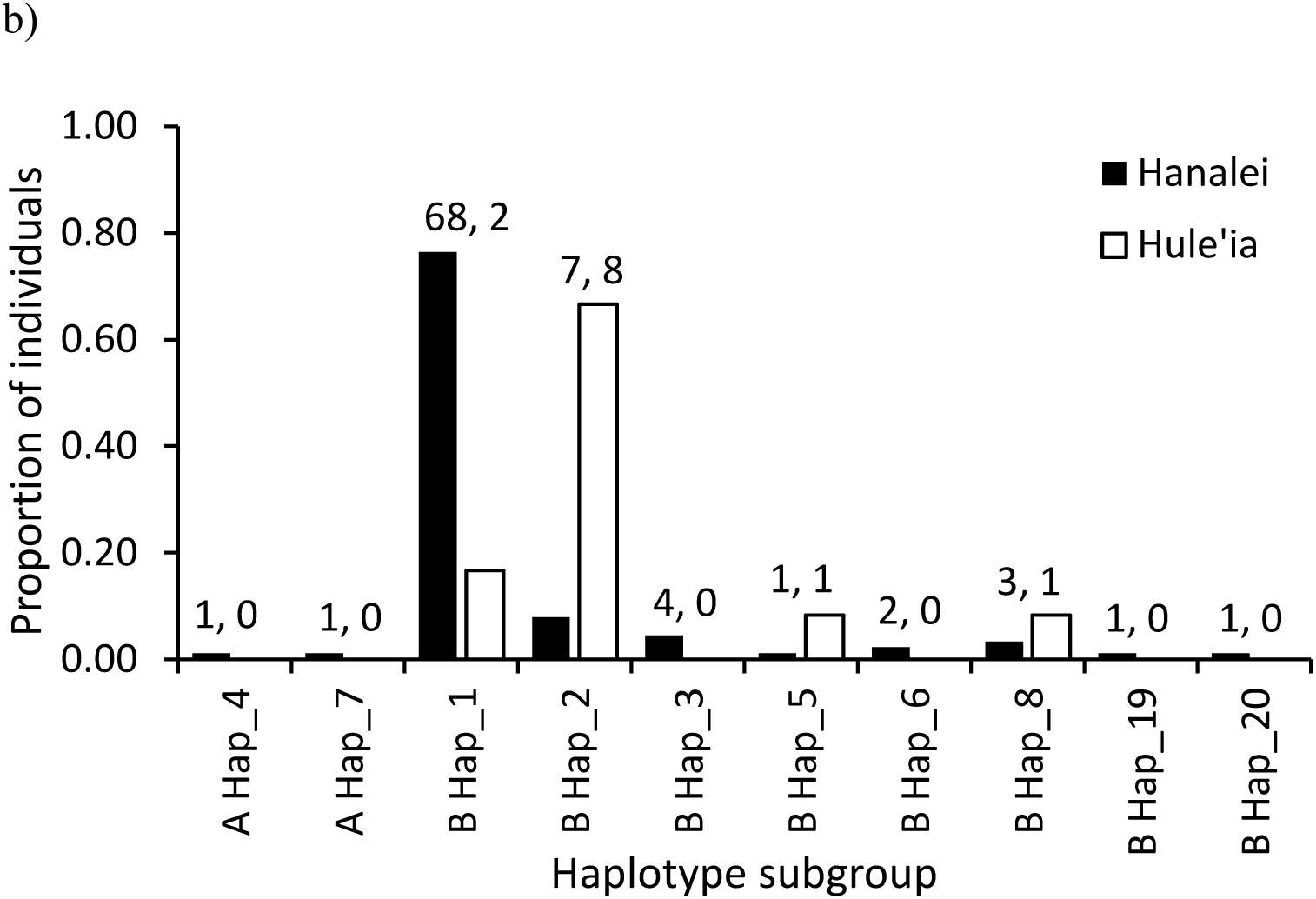
The proportions of different haplotypes that were found in a) adult males and adult females of koloa maoli sampled at Hanalei National Wildlife Refuge, Kaua‘i, Hawai‘i, USA and all sampled koloa on Hanalei and Hulē‘ia National Wildlife Refuges, Kaua‘i, Hawai‘i, USA. Koloa were sampled for haplotyping in 2011, and haplotyped koloa were captured from December 2010–November 2015. The sample sizes for each haplotype are provided above the bars, in order. A-type haplotypes (more common in mallards) were excluded for the G-test, and all haplotypes which were found in less than 4 individuals were combined into one category.

### Distance (meters)

#### Regional-scale relatedness and genetic structure

Among the subset of genotyped ducks that were captured as adults (*n*_Hanalei_ = 79; *n*_Hulē‘ia_ = 12), we found no closely related between-refuge dyads (Avg. ± SD: 0.0049 ± 0.0084; maximum *r* = 0.1071). In contrast, among adult dyads within refuges (Avg. ± SD: 0.0118 ± 0.0333), 34 dyads were closely related (r > 0.125, maximum *r* = 0.5660). Further, we found a significant difference in haplotype frequencies between the refuges (*G_adj_* = 19.3448, *df* = 4, *P* = 0.0007; Figure 7b), with the majority of sampled individuals in Hanalei having Haplotype 1 (corresponds to Haplotype J in Fowler *et al*. 2009), and the majority in Hulē‘ia having Haplotype 2 (corresponds to Haplotype N in Fowler *et al*. 2009).

## DISCUSSION

We present the first study on social associations and microspatial genetic structure in the endangered, island endemic koloa maoli. We found nonrandom associations that were not based on plumage traits, body mass, or relatedness. Further, we found some evidence for genetic structure in koloa between two core wetland sites on Kaua‘i, but no evidence for genetic structure within the largest site (Hanalei NWR). Our results have direct implications for koloa conservation: they suggest that koloa do not have a preference for associating with males with mallard-like phenotypes, that avian botulism outbreaks are unlikely to generate biased reductions in genetic diversity in the largest koloa population, and distinct mitochondrial diversity between refuges is important for a species that faces the risk of genetic extirpation. Further, we make some recommendations for future translocations based on our findings.

### Medium-term preferential associations among koloa

Koloa exhibited preferential associations among males and between males and females over extended periods. Being spatially restricted, experiencing high annual survival (Malachowski *et al*. 2022), and high local densities may have promoted repeated interactions between individual koloa, and, consequently, preferential associations. Although historically Hawai‘i had no native mammalian predators and few avian predators of native bird species (Burney *et al*. 2001), the introduction of rats (*Rattus* sp.), domestic cats (*Felis catus*), and pigs (*Sus scrofa*) has generated sometimes high levels of terrestrial predation on Hawaiian waterbirds (Malachowski 2020, Webber 2022), which may also promote grouping behavior. Associations in our study were defined as co-occurrence in a baited trap, and not any behavioral criteria, so observed trends could be a result of spatiotemporal overlap in habitat use. However, most of the koloa that were captured at least thrice and more than 30 days apart moved between units. The presence of preferential associations that last across months, therefore, suggests coordinated movement of associates. Lagged association rates suggest roughly seasonal (∼7 months) preferential associations among males, and annual (∼14 months) preferential associations between males and females (Supplementary Material Figure S6). The male-biased sex ratio might promote male associations, as males without a mate can associate with familiar conspecifics (see below for the benefits of foraging with familiar associates), as opposed to associating with a mate and competing with other males. Our results also suggest long-term pair bonds in koloa (also supported by the high percentage of pair bonded-females found throughout the year on Hanalei; see Malachowski *et al*. 2019), similar to the related island-dwelling Laysan duck (Reynolds *et al*. 2009) and other island ducks (Sorenson 1992). We cannot distinguish between social preferences and mate pair associations in our dataset. However, social associations before breeding may also influence reproductive success, by facilitating mate-pair formation (McDonald *et al*. 2020), sometimes years before their first breeding (Teitelbaum *et al*. 2017). However, there were a few instances of related (r > 0.125) male-female pairs caught in the same trap, so male-female associations might not exclusively be in the context of mating. Female-female associations were not different from random in this dataset, but fewer females in the refuge and lower capture rates of those females may have limited our ability to detect preferential associations. Alternatively, female koloa may truly be constrained in their ability to form longer-term associations with each other due to asynchronous periods of nest incubation and higher mortality.

### Plumage traits and body mass don’t seem to strongly influence associations

While hybrid zones in other birds showed assortative associations based on plumage characteristics (Zonana *et al*. 2019, Semenov *et al*. 2017), plumage traits and body mass did not play a detectable role in koloa associations. Our results suggest that, in koloa, mallard-like plumage is not selected for or against in social relationships. This could be due to their year-round breeding and monogamous mating system (Weller 1980, Malachowski *et al*. 2019), which is associated with less sexual dichromatism in ducks (Figuerola and Green 2000). Additionally, female mallards appear to select males based on bill coloration more than plumage (Omland 1996a, b); female koloa may also use pigmented soft parts and behavioral traits (e.g. courtship displays, defense, vigilance) instead of or in addition to plumage traits to select mates. In the case of body mass, as food is presumably available in the managed wetland habitat (USFWS 2020), there is possibly no pressure to associate with heavier individuals, who are likely successful foragers. We observed a moderate but insignificant negative relationship between female mass and their weighted degree in the two-year dataset; it is possible that heavier females are dominant, and avoided by lighter subordinate females as feeding associates to avoid aggression. As females often have different nutritional requirements (Reynolds and Perrins 2010), body mass might have a greater role to play in their sociality, and with more female captures, these patterns may become clearer. Instead of morphometric traits, it is possible that koloa form preferential association based on familiarity. A preference for familiar social associates has been observed in other birds (Kabasakal *et al*. 2017, Gomes *et al*. 2022), though males of some bird species preferred unfamiliar associates (Kohn *et al*. 2015). Foraging with familiar associates may provide direct benefits like low aggression levels and increased foraging and vigilance (Griffiths *et al*. 2004), and association with unrelated but familiar individuals may also be beneficial (Cameron *et al*. 2009, Riehl and Strong 2018).

### No spatial genetic structure within a refuge

Within Hanalei, koloa of both sexes move across units and genetic relatedness and mitochondrial haplotype did not strongly influence their associations or spatial distribution. However, Malachowski (2020) found higher regional fidelity for females than males, thus there may be sex bias in movement at slightly larger spatiotemporal scales. Although our spatial scale was small (inter-trap distances of ∼2-1500 meters), another island bird, the barnacle goose (*Branta leucopsis*), showed genetic structure due to philopatry at similar distances (see Anderholm *et al*. 2009). The management regime in Hanalei, where wetlands are managed seasonally or semi-annually, and taro cultivation is temporally staggered, results in spatiotemporal variability in habitat and food availability (USFWS 2020). This likely contributes to the observed movement and, possibly, also to the lack of strong spatial genetic structure within the refuge.

Even without kin discrimination (Leedale *et al*. 2020), the fact that we found few related (*r*>=0.125) male-female dyads captured together or near each other suggests low risk of inbreeding. The high number of unrelated associates and the movement across units by both sexes may allow birds in this population to find unrelated mates with ease, without showing spatial sub-structuring or strong sex-biased dispersal. However, the presence of some related male-female dyads in our dataset may hint at delayed dispersal in this species, as exhibited by other island birds (Covas 2012, Dibben-Young *et al*. 2019); examining juvenile dispersal in this population would yield more insight.

### Genetic structure between refuges

We found differences in mitochondrial haplotypes and no closely related dyads between Hanalei and Hulē‘ia wildlife refuges on Kaua‘i. Despite koloa being highly mobile and strong fliers (Engilis *et al*. 2002), these results support previous evidence of limited gene flow (Wells *et al*. 2019a) and movement (Malachowski 2020) between the two refuges. Though koloa can and do move across similar and greater distances than the distance between Hanalei and Hulē‘ia (>30 km; Malachowski 2020), dispersal between the refuges appears uncommon. Other island birds (Bertrand *et al*. 2014, Khimoun *et al*. 2017), including other waterfowl species (Anderholm *et al*. 2009, Fishman *et al*. 2011) show similar fine-scale genetic structure within an island.

### Implications for koloa conservation

The presence of a large number of conspecific males (and few feral mallards) and the male-biased sex ratio of the koloa population on Kaua’i has been suggested as the possible reason for the low levels of hybridization on the island (Wells *et al*. 2019a). Our results showing a lack of preference (or avoidance) for mallard-like plumage or heavier individuals (mallards are heavier than koloa; Weller 1980, Livezey 1993) lend further support this idea, although females could prefer mallard mates for other reasons such as behavior.

In this dataset, unrelated and distantly related dyads were caught in the same trap. Thus, during potential future translocations, moving individuals caught in the same trap together would both reduce the likelihood of disrupting preferential associations while also retaining genetic diversity among translocated individuals. Given the larger regional site fidelity in koloa (Malachowski 2020), fission-fusion dynamics (where koloa have a few preferred associates but also form many temporary and unstable associations) may enable translocated koloa to associate with residents of the translocated area (also see Kaczensky *et al*. 2021).

The lack of any strong spatial genetic structure within Hanalei means that, at the spatiotemporal scales relevant for avian botulism outbreaks, related koloa were not found closer to each other than unrelated koloa. Therefore, avian botulism outbreaks are unlikely to disproportionately remove kin clusters, which would bias reductions in genetic diversity in the largest koloa population. Further, as koloa didn’t seem to associate more with related than unrelated conspecifics, socially transmitted diseases like avian influenza (detected in koloa; Dugger *et al*. 2016) are also unlikely to selectively remove relatives and correlated genetic diversity. It is worth noting that avian botulism remains a primary threat and a year-round source of mortality to the population (Malachowski *et al*. 2022), and epizootics have the potential to affect other aspects of the population, like population dynamics and persistence (Perez-Heydrich *et al*. 2012). At a larger scale, there was evidence for genetic structure between the two largest refuges on the island; without connectivity via dispersal, local population recovery, if needed, through immigration is unlikely. Knowledge of such fine-scale population differentiation is important for management, and for maintaining a balance between retaining unique genetic diversity and overall demographic connectivity (Scribner *et al*. 2005).

In conclusion, we applied new analyses methods to existing datasets from a data-limited endangered species. Despite the limitations to the available data, we were able to shed light on aspects of their social behavior and spatial genetic structure, produce results that are of direct relevance to two of the biggest current threats to their conservation (avian botulism outbreaks and hybridization with mallards) and inform translocation strategy.

## ACKNOWLEDGEMENTS

The authors thank Kauaʻi NWR Complex (particularly S. N. Smith, M. W. Mitchell, C. C. Smith, and J. Campbell) for support. M. Reynolds, KUPU/Americorps, field assistants, and volunteers helped with sample collection. I. E. Engilis and undergraduate assistants of the UC Davis Museum of Wildlife and Fish Biology provided technical support for storing biological samples. The UC Davis Genomic Variation Lab and J. Peters provided lab space for original genetic work; J. DaCosta and M. Sorenson provided computational resources and advice during original genetic analyses. We thank C. Zagnoli-Robb for formatting preliminary data.

## FUNDING STATEMENT

This work was funded by Oregon State University, Colorado State University, USFWS, USGS, the Hawaiʻi Department of Land and Natural Resources, the Selma Herr Endowment, and the Dennis G. Raveling Endowment. This work used the Vincent J. Coates Genomics Sequencing

Laboratory at UC Berkeley, supported by NIH S10 Instrumentation Grants S10RR029668 and S10RR027303.

## ETHICS STATEMENT

The data collection was in compliance with the USFWS Threatened and Endangered Species Permit TE-702631 (Subpermits KNWR-7, KNWR-8, and KNWR-9), USFWS Region 1 Migratory Bird Depredation permit MB120267-1, Federal Bird Banding permit 23718, State of Hawai‘i Protected Wildlife permits WL12-03 and WL15-02, and Oregon State University Institutional Animal Care and Use protocols. Access to Hanalei NWR was granted by USFWS.

## CONFLICT OF INTEREST STATEMENT

All authors declare that they have no conflicts of interest.

## AUTHOR CONTRIBUTIONS

PK and CPW conceived the study and PK analyzed the data. CPM collected the field data and samples with help from BDD, KJU, AEJr, and CPW; CPW and PL generated and analyzed genetic data. PK and CPW wrote the initial draft, edited by CPM. All coauthors reviewed and edited subsequent drafts.

## DATA AVAILABILITY

Genetic data are available (ddRAD:SRA BioProject PRJNA577581, Sample accession IDs SAMN13029772–SAMN13030226; mtDNA: GenBank Accession Num. MN563303–MN563571 and MN603671–MN603691, other data files: Dryad accession https://doi.org/10.5061/dryad.ffbg79cqf (Wells *et al*., 2019b)). Association data files will be archived on Dryad upon acceptance.

## Supplementary Material

**Figure S1:**
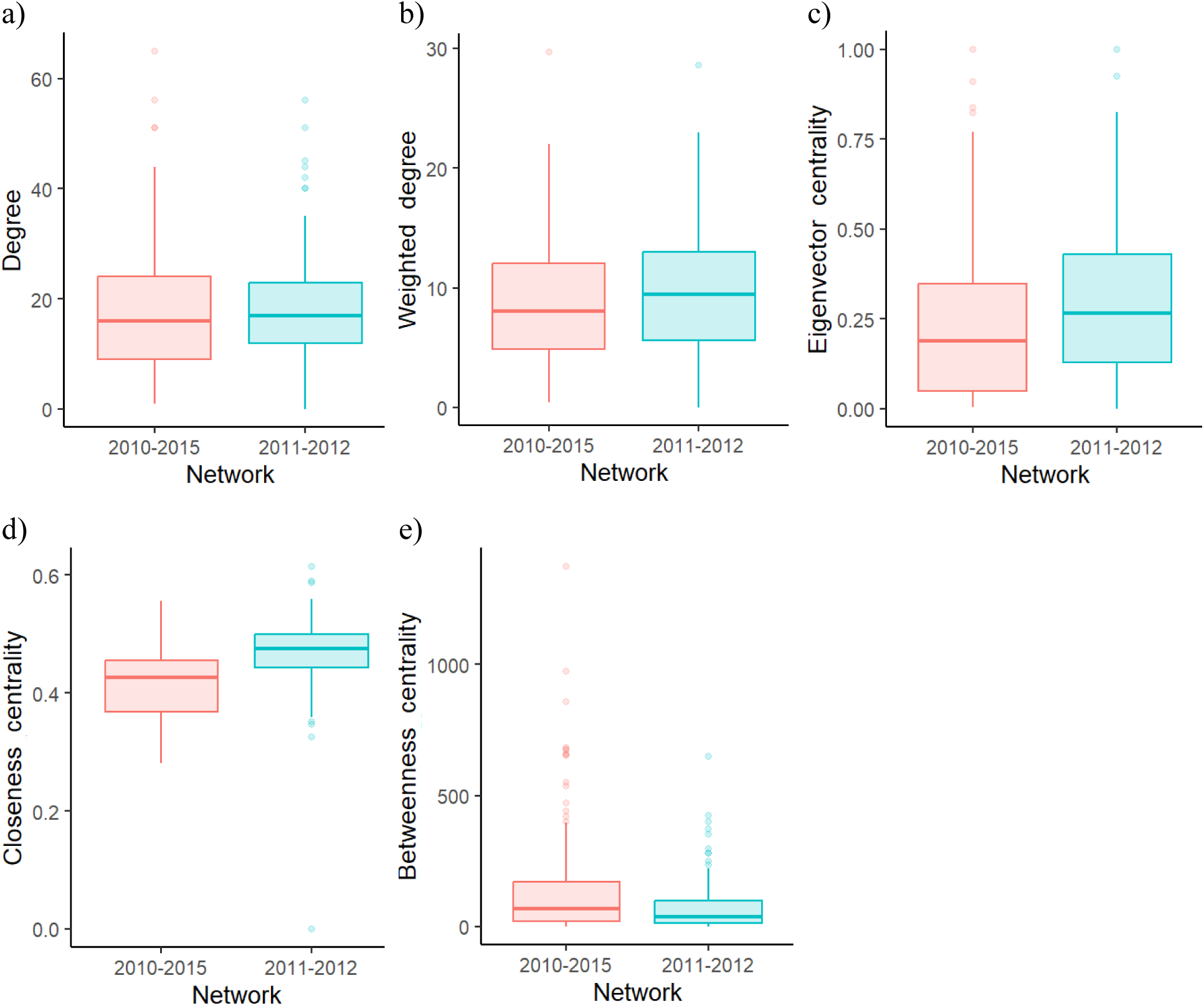
Network metrics using the overall data and the two-year dataset. We created social networks of koloa (sighted at least thrice) associations, using all captures (2010-2015), and using a subset of the captures in two years (2011-2012). Network metrics (calculated using Gephi 0.10; Bastian *et al*. 2009) of the networks overlapped, despite differences in the number of nodes (190 koloa in the overall dataset, 133 koloa in the two-year dataset). Distribution of a) degree (the number of associates of a node), b) weighted degree (the strength of associations or the sum of association indices of a node with all other nodes), c) eigenvector centrality (a measure of the influence of the node; a node with high eigenvector centrality tends to be connected to other nodes with high eigenvector centrality), d) closeness centrality (reciprocal of the sum of shortest path lengths from node to all other nodes), and e) betweenness centrality (the number of shortest paths, between all pairs of nodes in the network, that pass through a node) values in the overall network (2010-2015; in pink and to the left) and the two-year network (2011-2012; in blue and to the right).

**Figure S2:**
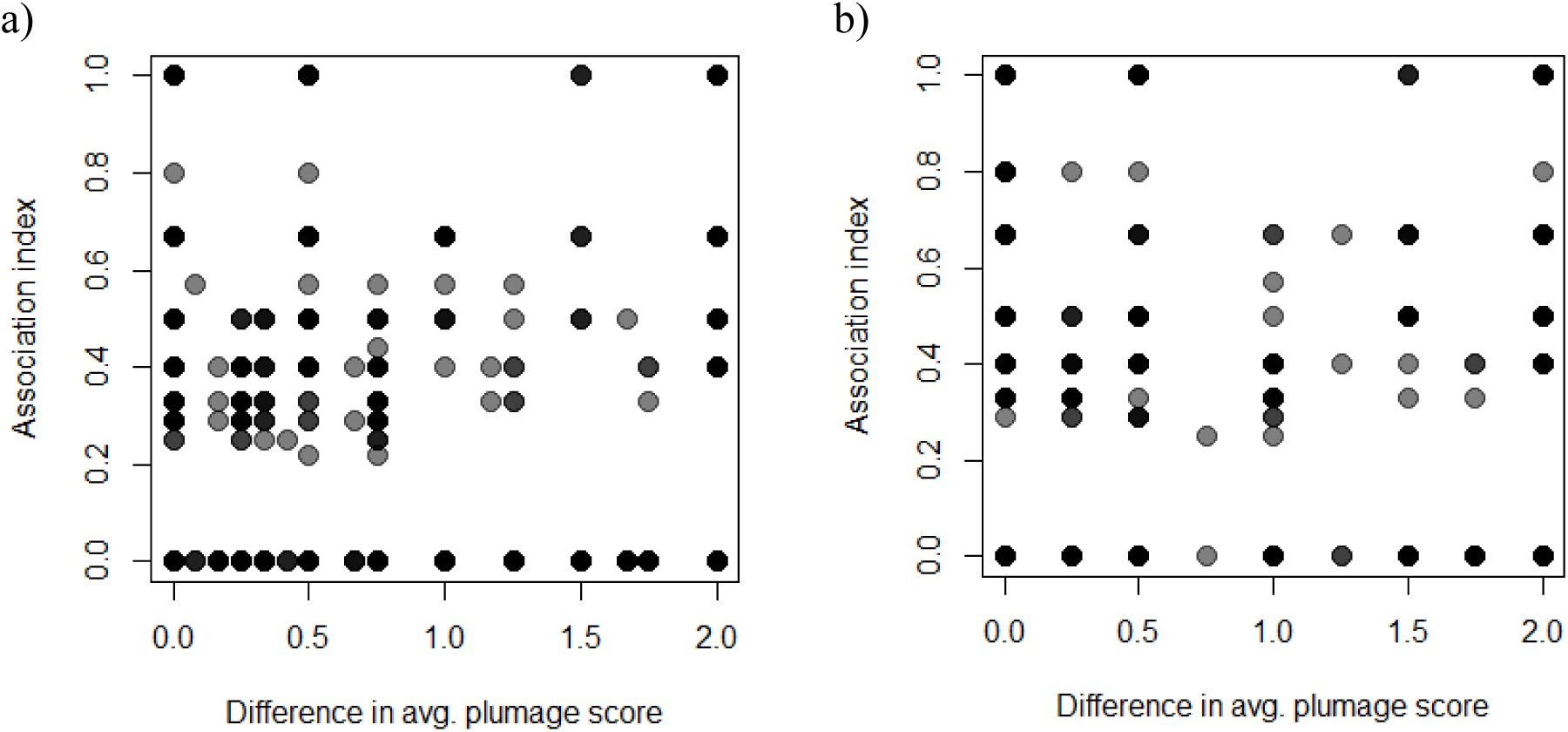
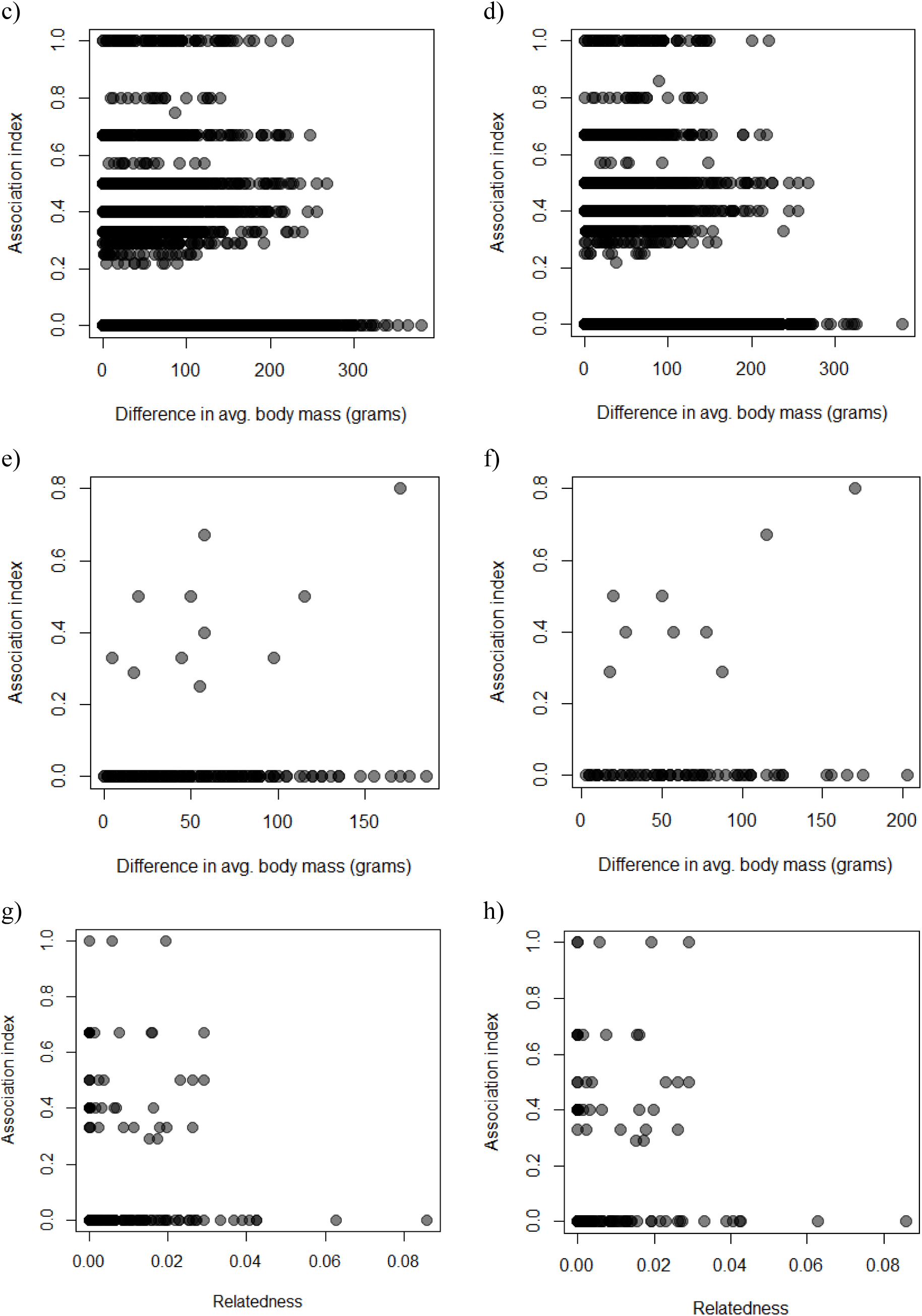
Scatter plots of association index values and various traits. Association index values and differences in average plumage score of male-male dyads a) using 2010-2015 captures and b) using 2011-2012 captures, differences in average body mass of male-male dyads a) c) using 2010-2015 captures and d) using 2011-2012 captures, differences in average body mass of female-female dyads e) using 2010-2015 captures and f) using 2011-2012 captures, and relatedness and association index of male-male dyads g) using 2010-2015 captures and h) using 2011-2012 captures. Only koloas sighted 3 or more times in the corresponding dataset were used for all plots.

**Figure S3.**
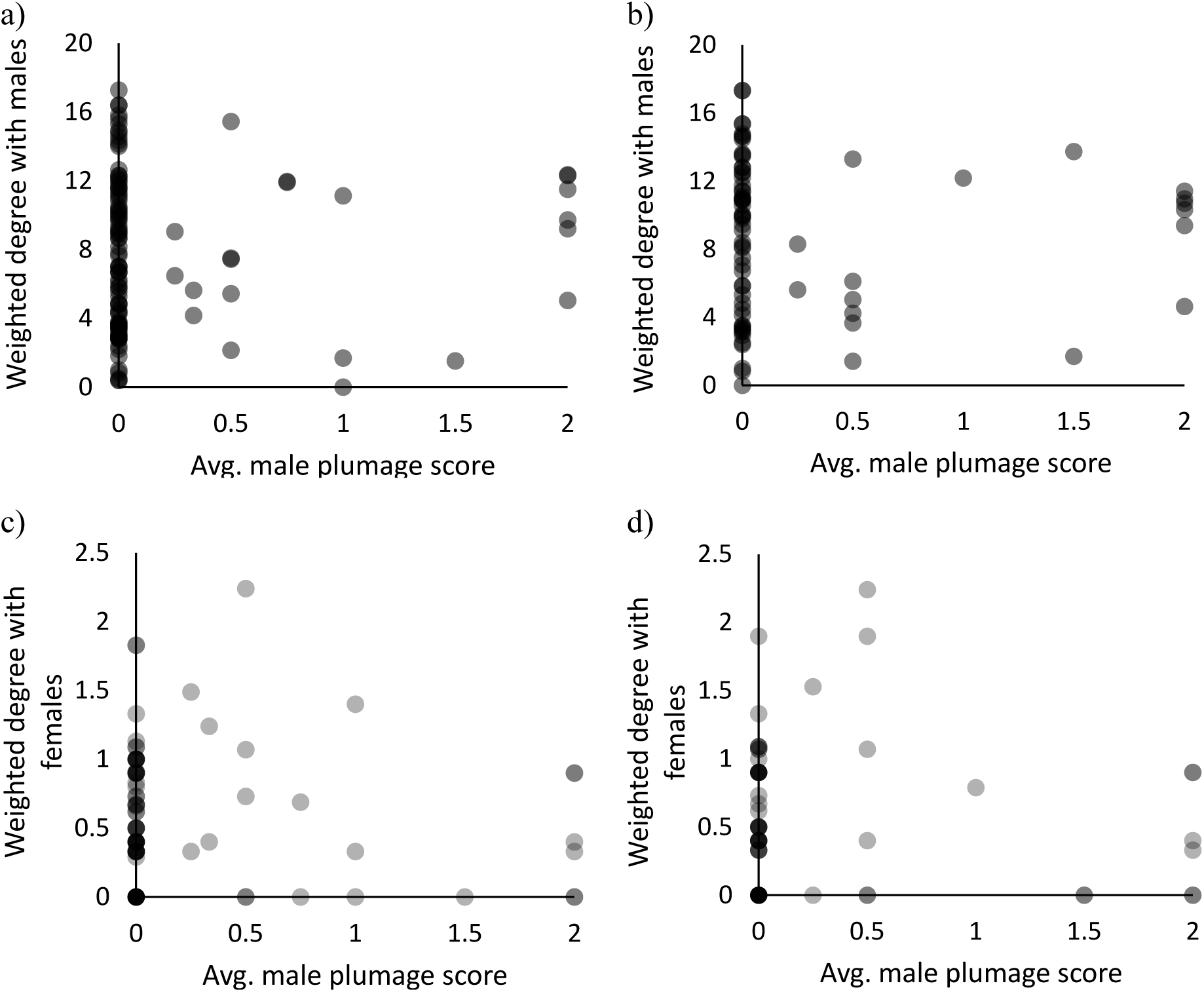
Scatter plots of the average plumage score of males and their weighted degree. (or association strength) with other males in the network with 2010-2015 captures (a) and with 2011-2012 captures (b), and their weighted degree with females in the network with 2010-2015 captures (c) and with 2011-2012 captures (d). Captures from November-April were used for all plots with plumage score. Only koloas sighted 3 or more times in the corresponding dataset were used for all plots.

**Figure S4.**
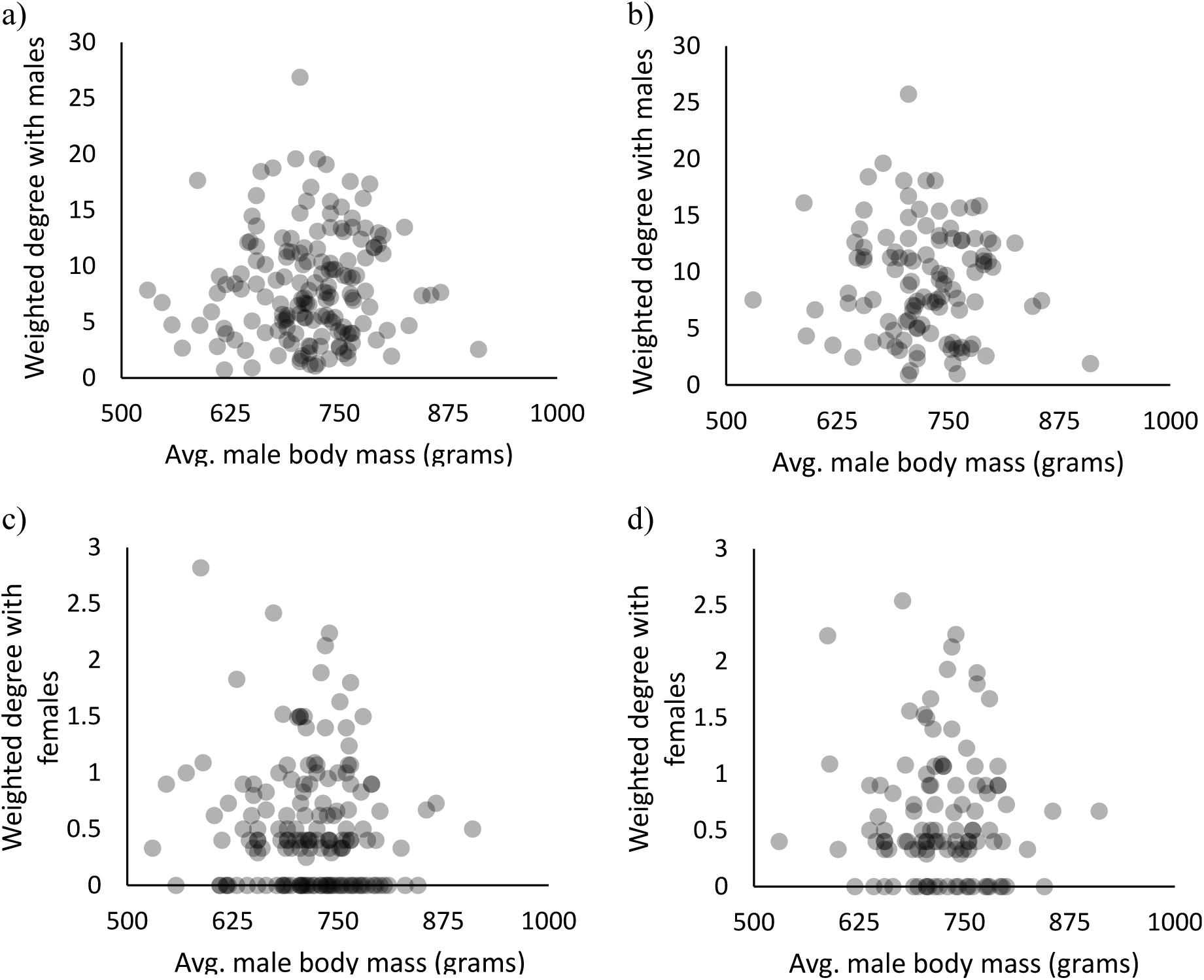
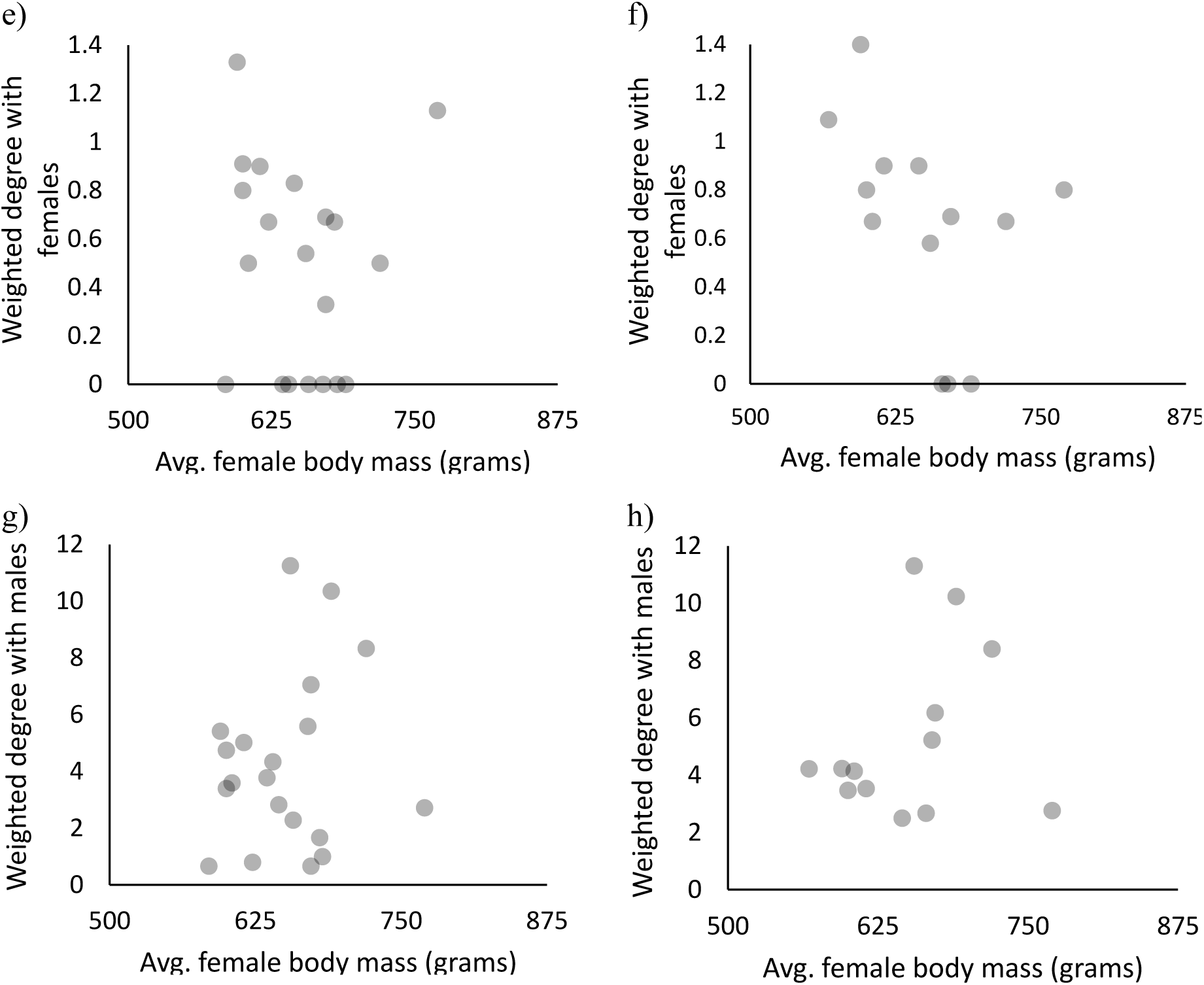
**Scatter plots of the average body mass of males and their weighted degree** with other males in the network with 2010-2015 captures (a) and with 2011-2012 captures (b), and their weighted degree with females in the network with 2010-2015 captures (c) and with 2011- 2012 captures (d). **Scatter plots of the average body mass of females and their weighted degree** with other females in in the network with 2010-2015 captures (e) and with 2011-2012 captures (f), and their weighted degree with males in the network with 2010-2015 captures (g) and with 2011-2012 captures (h). Only koloas sighted 3 or more times in the corresponding dataset were used for all plots.

**Figure S5.**
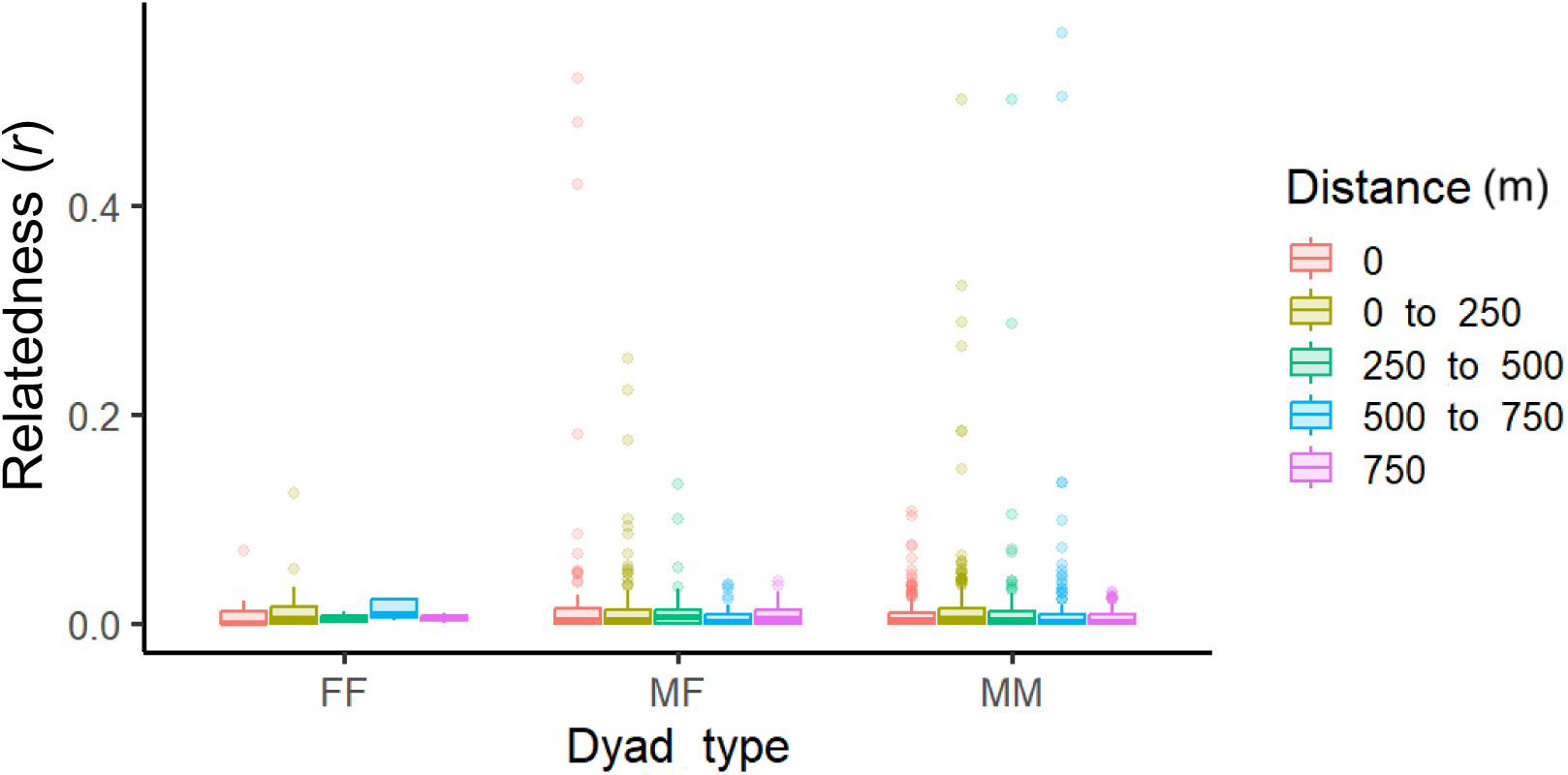
*r* (relatedness) values binned by distance (in meters) of female-female (FF), male-female (MF) and male-male (MM) dyads.

**Figure S6:**
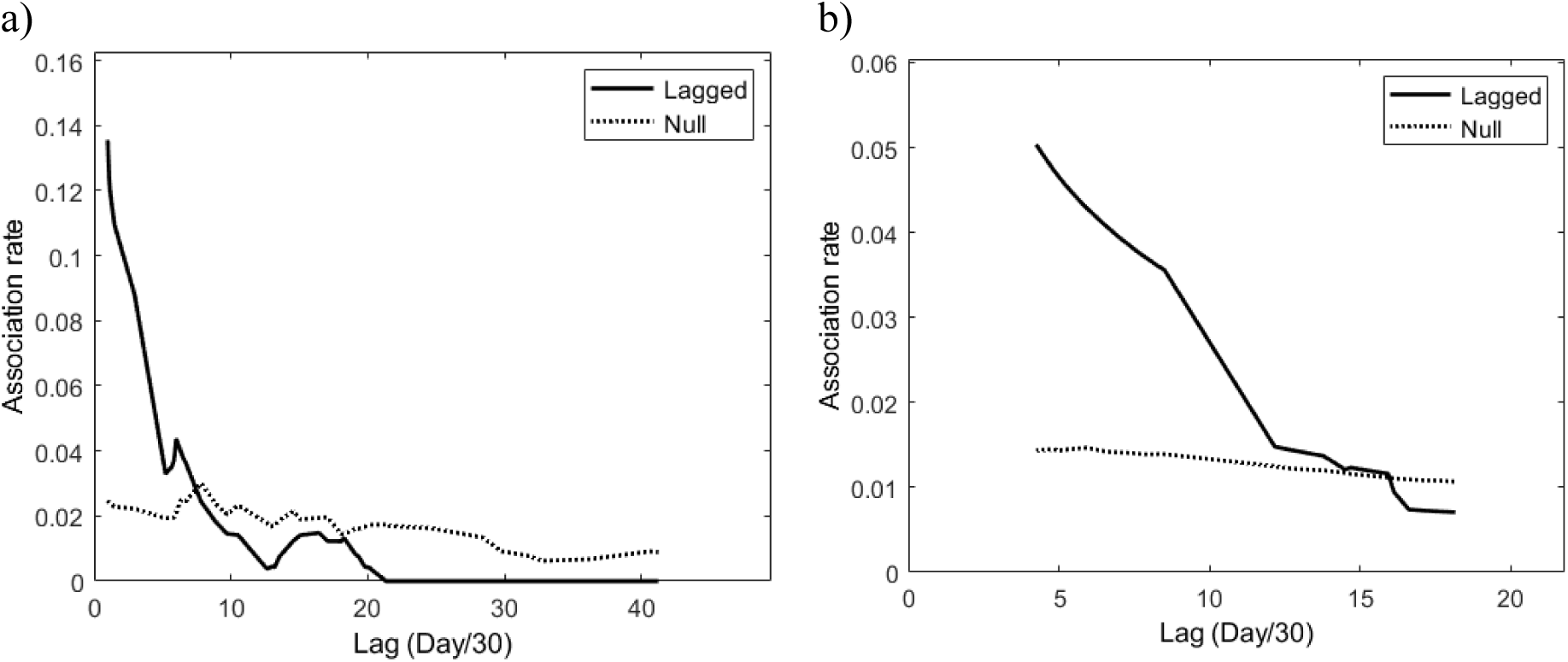
Lagged associations rates for male-male and male-female associations. For dyads that showed preferential associations, we evaluated temporal patterns of those associations by calculating the lagged association rates (LAR; i.e., the probability that two individuals that are associated at a given time will be associated again after various time lags; Whitehead 1995). We compared LAR with null association rates (NAR; i.e., expected values if individuals associated at random) for different time lags (in our case, multiples of the 30-day sampling period). Preferential associations last as long as LAR are greater than NAR. Male-male preferential associations (a) seem to last for ∼7 sampling periods, while male-female preferential associations (b) last for ∼14 sampling periods.

**Table S1:** Number of trap-days during which we attempted to capture koloa maoli using baited swim-in traps during December 2010–November 2015 at Hanalei National Wildlife Refuge, Kaua‘i, Hawai‘i USA. We used trapping data from those traps where the identities of all trapped ducks in one trap were clear. The years 2011 and 2012 were most intensely sampled (both in terms of days sampled, and the number of months sampled), and we repeated some of the network analyses using this two-year dataset.

| Year | Jan | Feb | Mar | Apr | May | Jun | Jul | Aug | Sep | Oct | Nov | Dec |
| --- | --- | --- | --- | --- | --- | --- | --- | --- | --- | --- | --- | --- |
| 2010 | 0 | 0 | 0 | 0 | 0 | 0 | 0 | 0 | 0 | 0 | 0 | 19 |
| 2011 | 11 | 11 | 8 | 5 | 0 | 10 | 1 | 9 | 0 | 0 | 0 | 0 |
| 2012 | 0 | 0 | 0 | 0 | 13 | 9 | 0 | 0 | 0 | 0 | 8 | 12 |
| 2013 | 0 | 0 | 0 | 0 | 0 | 0 | 0 | 0 | 0 | 0 | 2 | 19 |
| 2014 | 0 | 0 | 0 | 0 | 0 | 5 | 0 | 0 | 0 | 0 | 0 | 0 |
| 2015 | 0 | 0 | 0 | 0 | 0 | 0 | 0 | 0 | 0 | 0 | 9 | 0 |

**Table S2.** Scoring scheme for plumage assessment of male koloa. While the scores were assigned as a binary (0 or 1), there is a range of plumage variation expressed by genetically vetted koloa.

| Trait | Mallard-like plumage | Score | Not mallard-like plumage | Score |
| --- | --- | --- | --- | --- |
| % Green on Head | >50% | 1 | <50% | 0 |
| Chest Color/Pattern | Chestnut/Unmarked | 1 | Marked | 0 |
| Belly and Flank Pattern | Unmarked | 1 | Marked | 0 |
| Back Pattern | Unmarked | 1 | Marked | 0 |

**Table S3:** Choosing a relatedness estimator. We ran three simulations using the empirical allele frequencies of 1502 non-homozygous loci with no missing data. As these loci do not have any missing data, the missing rate was kept as 0 for all three simulations. Simulation 1 was run with 0 error and dropout rates, Simulation 2 with 0.01 error and dropout rates, and Simulation 3 with 0.05 error and dropout rates. Relatedness was calculated between simulated genotypes of 100 parent-offspring (*r*=0.5), 100 full siblings (*r*=0.5), 100 half siblings (*r*=0.25), 100 first cousins (*r*=0.125), and 100 unrelated (*r*=0.0) dyads using all seven estimators. In each simulation, for each estimator, the Pearson’s correlation between the true relatedness and the estimated relatedness was calculated. Further, as one of the 108 koloa was sampled twice, we included both samples in the analysis, and estimated coefficients of relatedness between the 109 samples using all seven estimators. We checked whether the estimators calculated the relationship between the identical pair accurately (*r*=1.0). While all estimators, across the three simulations, calculated relatedness estimates that were close to the real value, the DyadML estimator consistently had the highest Pearson’s correlation coefficient (value in bold). DyadML also estimated the r value (1.0) between two samples of the same individual. We re-ran the relatedness estimate analysis for the 108 unique samples (as the inclusion of the duplicate sample would have modified the allele frequencies and the relatedness estimates) in Coancestry. The DyadML relatedness estimates from this run were used for further analyses.

| Simulation | Trio ML | Wang | Lynch Li | Lynch<br>Ritland | Ritland | Queller<br>Goodnight | Dyad<br>ML |
| --- | --- | --- | --- | --- | --- | --- | --- |
| 1 | 0.995661 | 0.993375 | 0.992963 | 0.986643 | 0.963747 | 0.993354 | **0.995693** |
| 2 | 0.996041 | 0.993325 | 0.992826 | 0.988876 | 0.964984 | 0.993477 | **0.996053** |
| 3 | 0.993803 | 0.990509 | 0.989772 | 0.986026 | 0.961561 | 0.990669 | **0.994054** |

**Table S4:** Testing for preferential associations. We used the half-weight index (HWI) and the ‘*Permute associations within samples*’ method to look for preferential associations among males, using a 30-day sampling period. This permutation method controls for individual differences in gregariousness. We found that the CV of the real dataset was significantly greater than the permuted datasets (recommended test for long-term preferential associations). We also used the social affinity index (SAI) and the ‘*Permute groups within samples*’ method to check for preferential associations between males and females, and again found evidence for long-term preferential associations (CV of the real dataset was significantly greater than the permuted datasets; Table 1). In both cases, number of permutations were increased from 10,000 to 20,000, and *P*-values remained similar. Results for 20,000 permutations are reported. *P*= 1 - (No. of permutations where the real dataset’s CV is >Random data’s CV/20000). Significant *P*-values are highlighted in bold. associations within samples within samples

| Permutation | CV<sub>observed</sub> | Mean of CV<sub>permuted</sub> | *P* |
| --- | --- | --- | --- |
| Male-Male associations HWI, permute associations within samples | 7.1258 | 7.1052 | **<0.0001** |
| Male-Female associations SAI, permute groups within samples | 8.7001 | 8.6212 | **0.0003** |

## Notes

### Competing Interest Statement

The authors have declared no competing interest.

## References

Anderholm, S., P. Waldeck, H. P. Van der Jeugd, R. C. Marshall, K. Larsson, and M. Andersson (2009). Colony kin structure and host-parasite relatedness in the barnacle goose. Molecular Ecology 18(23): 4955–4963.

Banko, W. E. (1987). Historical synthesis of recent endemic Hawaiian birds. Koloa-Maoli. Manoa, HI: University of Hawaii.

Bastian, M., S. Heymann, and M. Jacomy (2009). Gephi: an open source software for exploring and manipulating networks. In Third international AAAI conference on weblogs and social media.

Beauchamp, G. (2004). Reduced flocking by birds on islands with relaxed predation. Proceedings of the Royal Society of London. Series B: Biological Sciences 271(1543): 1039–1042.

Beauchamp, G. (2010). Relaxed predation risk reduces but does not eliminate sociality in birds. Biology Letters 6(4): 472–474.

Beck, K. B., C. E. Regan, K. McMahon, S. Crofts, E. F. Cole, J. A. Firth, and B. C. Sheldon (2024). Experimental manipulation of population density in a wild bird alters social structure but not patch discovery rate. Animal Behaviour 209: 95–120.

Bertrand, J. A. M., Y. X. C. Bourgeois, B. Delahaie, T. Duval, R. García-Jiménez, J. Cornuault, P. Heeb, B. Milá, B. Pujol, and C. Thébaud (2014). Extremely reduced dispersal and gene flow in an island bird. Heredity 112(2): 190–196.

Blondel, J. (2000). Evolution and ecology of birds on islands: trends and prospects. Vie et Milieu/Life & Environment 205–220.

Brooks, M. E., K. Kristensen, K. J. Van Benthem, A. Magnusson, C. W. Berg, A. Nielsen, H. J. Skaug, M. Mächler, and B. M. Bolker (2017). glmmTMB balances speed and flexibility among packages for zero-inflated generalized linear mixed modeling. The R journal 9(2): 378–400.

Burney, D. A., H. F. James, L. P. Burney, S. L. Olson, W. Kikuchi, W. L. Wagner, M. Burney, D. McCloskey, D. Kikuchi, F. V. Grady, and R. Gage (2001). Fossil evidence for a diverse biota from Kaua ‘i and its transformation since human arrival. Ecological Monographs 71(4): 615–641.

Cameron, E. Z., T. H. Setsaas, and W. L. Linklater (2009). Social bonds between unrelated females increase reproductive success in feral horses. Proceedings of the National Academy of Sciences 106(33): 13850–13853.

Carter, N. H., E. C. Wilson, and B. Gurung (2023). Social networks of solitary carnivores: The case of endangered tigers and insights on their conservation. Conservation Science and Practice 5(8): e12976.

Cheek, R. G., B. R. Forester, P. E. Salerno, D. R. Trumbo, K. M. Langin, N. Chen, T. S. Sillett, S. A. Morrison, C. K. Ghalambor, and W. C. Funk (2022). Habitat-linked genetic variation supports microgeographic adaptive divergence in an Island-endemic bird species. Molecular Ecology 31(10): 2830–2846.

Covas, R. (2012). Evolution of reproductive life histories in island birds worldwide. Proceedings of the Royal Society B: Biological Sciences 279(1733): 1531–1537.

Dibben-Young, A., K. C. Harmon, A. Lunow-Luke, J. L. Idle, D. L. Christensen, and M. R. Price (2021). Cooperative breeding behaviors in the Hawaiian Stilt (*Himantopus mexicanus knudseni*). Ecology and Evolution 11(10): 5010–5016.

Duffy, G. A., T. W. Pike, and K. N. Laland (2009). Size-dependent directed social learning in nine-spined sticklebacks. Animal Behaviour 78(2): 371–375.

Dugger, B., C. Malachowski, D. Heard, and K. Uyehara (2016). A Health Survey for Hawaiian Duck (*Anas wyvilliana*) at Hanalei National Wildlife Refuge, Kaua ‘i.

Engilis, A., K. J. Uyehara, and J. G. Giffin (2002). Hawaiian Duck: Anas wyvilliana. Birds of North America, Incorporated. https://birdsoftheworld.org/bow/species/hawduc/cur/introduction

Farine, D. R. (2017). A guide to null models for animal social network analysis. Methods in Ecology and Evolution 8(10): 1309–1320.

Figuerola, J., and A. J. Green (2000). The evolution of sexual dimorphism in relation to mating patterns, cavity nesting, insularity and sympatry in the Anseriformes. Functional Ecology 14(6): 701–710.

Fishman, D. J., S. R. Craik, D. Zadworny, and R. D. Titman (2011). Spatial-genetic structuring in a red-breasted merganser (Mergus serrator) colony in the Canadian Maritimes. Ecology and Evolution 1(2): 107–118.

Fowler, A. C., J. M. Eadie, and A. Engilis (2009). Identification of endangered Hawaiian ducks (*Anas wyvilliana*), introduced North American mallards (*A. platyrhynchos*) and their hybrids using multilocus genotypes. Conservation Genetics 10: 1747–1758.

Fromm, A., and S. Meiri (2021). Big, flightless, insular and dead: Characterising the extinct birds of the Quaternary. Journal of Biogeography 48(9): 2350–2359.

Fukunaga, K., C. P. Wells, and Lavretsky, P. (2026). Determining sex ratios and mitochondrial haplotypes of Hawai ‘i’s endemic and introduced ducks. Journal of Ornithology 167: 211–222.

Goldenberg, S. Z., M. A. Owen, J. L. Brown, G. Wittemyer, Z. M. Oo, and P. Leimgruber (2019). Increasing conservation translocation success by building social functionality in released populations. Global Ecology and Conservation 18: e00604.

Gomes, A. C. R., P. Beltrão, N. J. Boogert, and G. C. Cardoso (2022). Familiarity, dominance, sex and season shape common waxbill social networks. Behavioral Ecology 33(3): 526–540.

Goslee, S. C., and D. L. Urban (2007). The ecodist package for dissimilarity-based analysis of ecological data. Journal of Statistical Software 22: 1–19.

Grear, D. A., M. D. Samuel, K. T. Scribner, B. V. Weckworth, and J. A. Langenberg (2010). Influence of genetic relatedness and spatial proximity on chronic wasting disease infection among female white-tailed deer. Journal of Applied Ecology 47(3): 532–540.

Griffiths, S. W., S. Brockmark, J. Höjesjö, and J. I. Johnsson (2004). Coping with divided attention: the advantage of familiarity. Proceedings of the Royal Society of London. Series B: Biological Sciences 271(1540): 695–699.

Hartig, F. (2018). DHARMa: residual diagnostics for hierarchical (multi-level/mixed) regression models. R Packag version 020.

Jezierski, M. T., W. J. Smith, and S. M. Clegg (2023). The island syndrome in birds. Journal of Biogeography 51(9): 1607–1622.

Kabasakal, B., M. Poláček, A. Aslan, H. Hoi, A. Erdoğan, and M. Griggio (2017). Sexual and non-sexual social preferences in male and female white-eyed bulbuls. Scientific Reports 7(1): 5847.

Kaczensky, P., A. Salemgareyev, J. D. Linnell, S. Zuther, C. Walzer, N. Huber, and T. Petit (2021). Post-release movement behaviour and survival of Kulan reintroduced to the steppes and deserts of central Kazakhstan. Frontiers in Conservation Science, 2, 703358.

Khimoun, A., W. Peterman, C. Eraud, B. Faivre, N. Navarro, and S. Garnier (2017). Landscape genetic analyses reveal fine-scale effects of forest fragmentation in an insular tropical bird. Molecular Ecology 26(19): 4906–4919.

Kohn, G. M., G. R. Meredith, F. R. Magdaleno, A. P. King, and M. J. West (2015). Sex differences in familiarity preferences within fission–fusion brown-headed cowbird, Molothrus ater, flocks. Animal Behaviour 106: 137–143.

Langin, K. M., T. S. Sillett, W. C. Funk, S. A. Morrison, M. A. Desrosiers, and C. K. Ghalambor (2015). Islands within an island: repeated adaptive divergence in a single population. Evolution 69(3): 653–665.

Lavretsky, P., A. Engilis Jr, J. M. Eadie, and J. L. Peters (2015). Genetic admixture supports an ancient hybrid origin of the endangered Hawaiian duck. Journal of Evolutionary Biology 28(5): 1005–1015.

Leedale, A. E., J. Li, and B. J. Hatchwell (2020). Kith or kin? Familiarity as a cue to kinship in social birds. Frontiers in Ecology and Evolution 8: 77.

Livezey, B. C. (1993). Comparative morphometrics of Anas ducks, with particular reference to the Hawaiian Duck *Anas wyvilliana*, Laysan Duck A. laysanensis, and Eaton’s Pintail A. eatoni. Wildfowl 44(44): 75–100.

Madden, J. R., J. A. Drewe, G. P. Pearce, and T. H. Clutton-Brock (2011). The social network structure of a wild meerkat population: 3. Position of individuals within networks. Behavioral Ecology and Sociobiology 65(10): 1857–1871.

Malachowski, C. P. (2013). Hawaiian duck (Anas wyvilliana) behavior and response to wetland habitat management at Hanalei National Wildlife Refuge on Kaua’i. Master’s dissertation, Oregon State University, Corvallis, USA.

Malachowski, C. P. (2020). Movement ecology and population dynamics of the endangered Hawaiian Duck (Anas wyvilliana). Ph.D. dissertation, Oregon State University, Corvallis, USA.

Malachowski, C. P., and B. D. Dugger (2018). Hawaiian duck behavioral patterns in seasonal wetlands and cultivated taro. The Journal of Wildlife Management 82(4): 840–849.

Malachowski, C. P., B. D. Dugger, and K. J. Uyehara (2019). Seasonality of life history events and behavior patterns in the island endemic Hawaiian Duck (*Anas wyvilliana*). Waterbirds 42(1): 78–89.

Malachowski, C. P., B. D. Dugger, K. J. Uyehara, and M. H. Reynolds (2018). Nesting ecology of the Hawaiian Duck *Anas wyvilliana* on northern Kaua ‘i, Hawai ‘i, USA. Wildfowl 68(68): 123–139.

Malachowski, C. P., B. D. Dugger, K. J. Uyehara, and M. H. Reynolds (2022). Avian botulism is a primary, year-round threat to adult survival in the endangered Hawaiian Duck on Kaua ‘i, Hawai ‘i, USA. Ornithological Applications 124(2): duac007.

Mantel, N. (1967). The detection of disease clustering and a generalized regression approach. Cancer research 27: 209–220.

Matthews, T. J., J. P. Wayman, P. Cardoso, F. Sayol, J. P. Hume, W. Ulrich, J. A. Tobias, F. C. Soares, C. Thébaud, T. E. Martin, and K. A. Triantis (2022). Threatened and extinct island endemic birds of the world: Distribution, threats and functional diversity. Journal of Biogeography 49(11): 1920–1940.

McDonald, G. C., N. Engel, S. S. Ratão, T. Székely, and A. Kosztolányi (2020). The impact of social structure on breeding strategies in an island bird. Scientific Reports 10(1): 13872.

Morris, S. D., B. W. Brook, K. E. Moseby, and C. N. Johnson (2021). Factors affecting success of conservation translocations of terrestrial vertebrates: a global systematic review. Global Ecology and Conservation 28: e01630.

Omland, K. E. (1996a). Female mallard mating preferences for multiple male ornaments: I. Natural variation. Behavioral Ecology and Sociobiology 39: 353–360.

Omland, K. E. (1996b). Female Mallard mating preferences for multiple male ornaments: II. Experimental variation. Behavioral Ecology and Sociobiology 39: 361–366.

Paxton, E. H., K. Brinck, A. Henry, A. Siddiqi, R. Rounds, and J. Chutz (2021). Distribution and trends of endemic Hawaiian waterbirds. Waterbirds 44(4): 425–437.

Perez-Heydrich, C., M. K. Oli, and M. B. Brown (2012). Population-level influence of a recurring disease on a long-lived wildlife host. Oikos 121(3): 377–388.

Reynolds, M. H., J. H. Breeden, M. S. Vekasy, and T. M. Ellis (2009). Long-term pair bonds in the Laysan Duck. The Wilson Journal of Ornithology 121(1): 187–190.

Reynolds, S. J., and C. M. Perrins (2010). Dietary calcium availability and reproduction in birds. Current Ornithology Volume 17: 31–74.

Riehl, C., and M. J. Strong (2018). Stable social relationships between unrelated females increase individual fitness in a cooperative bird. Proceedings of the Royal Society B: Biological Sciences 285(1876): 20180130.

Rocke, T. E., and T. K. Bollinger (2007). Avian botulism. In Infectious Diseases of Wild Birds (N. J. Thomas, D. B. Hunter and C. T. Atkinson, Editors). Blackwell Publishing, Ames, IA, USA.

Ruckstuhl, K. E., and P. Neuhaus (2002). Sexual segregation in ungulates: a comparative test of three hypotheses. Biological Reviews 77(1): 77–96.

Scribner, K. T., J. A. Blanchong, D. J. Bruggeman, B. K. Epperson, C. Y. Lee, Y. W. Pan, R. I. Shorey, H. H. Prince, S. R. Winterstein, and D. R. Luukkonen (2005). Geographical genetics: conceptual foundations and empirical applications of spatial genetic data in wildlife management. The Journal of wildlife management 69(4): 1434–1453.

Semenov, G. A., E. S. Scordato, D. R. Khaydarov, C. C. Smith, N. C. Kane, and R. J. Safran (2017). Effects of assortative mate choice on the genomic and morphological structure of a hybrid zone between two bird subspecies. Molecular Ecology 26(22): 6430–6444.

Silk, M. J., R. A. McDonald, R. J. Delahay, D. Padfield, and D. J. Hodgson (2021). CMRnet: An r package to derive networks of social interactions and movement from mark–recapture data. Methods in Ecology and Evolution 12(1): 70–75.

Snijders, L., D. T. Blumstein, C. R. Stanley, and D. W. Franks (2017). Animal social network theory can help wildlife conservation. Trends in ecology & evolution 32(8): 567–577.

Sokal, R. R., and F. J. Rohlf (1981). Biometry WH Freeman and Company. New York, 14.

Sonsthagen, S. A., S. L. Talbot, R. B. Lanctot, and K. G. McCracken (2010). Do common eiders nest in kin groups? Microgeographic genetic structure in a philopatric sea duck. Molecular Ecology 19(4): 647–657.

Sorenson, L. G. (1992). Variable mating system of a sedentary tropical duck: the white-cheeked pintail (*Anas bahamensis bahamensis*). The Auk 109(2): 277–292.

Taylor, L. U., E. Benavides, J. W. Simmons, and T. J. Near (2021). Genomic and phenotypic divergence informs translocation strategies for an endangered freshwater fish. Molecular Ecology 30(14): 3394–3407.

Teitelbaum, C. S., S. J. Converse, and T. Mueller (2017). Birds choose long-term partners years before breeding. Animal Behaviour 134: 147–154.

U.S. Fish and Wildlife Service. 2011 (USFWS). Recovery Plan for Hawaiian Waterbirds, Second Revision. U.S. Fish and Wildlife Service, Portland, Oregon.

U.S. Fish and Wildlife Service. 2020 (USFWS). Draft wetlands management and waterbird conservation plan. Hanalei National Wildlife Refuge. U.S. Fish and Wildlife Service, Portland, Oregon.

Wang, J. (2011). COANCESTRY: a program for simulating, estimating and analysing relatedness and inbreeding coefficients. Molecular ecology resources 11(1): 141–145.

Webber, B. M. (2022). Reproductive Success of the Hawaiian Common Gallinule (Gallinula galeata sandvicensis) Nesting in Taro and Managed Wetlands on Kaua‘i, Hawai‘i. Master’s dissertation, Oregon State University, Corvallis, USA.

Weiss, M. N., D. W. Franks, L. J. Brent, S. Ellis, M. J. Silk, and D. P. Croft (2021). Common datastream permutations of animal social network data are not appropriate for hypothesis testing using regression models. Methods in Ecology and Evolution 12(2): 255–265.

Weller M.W. (1980). The island waterfowl. Iowa State University Press.

Wells, C. P., P. Lavretsky, M. D. Sorenson, J. L. Peters, J. M. DaCosta, S. Turnbull, K. J. Uyehara, C. P. Malachowski, B. D. Dugger, J. M. Eadie, and A. Engilis Jr (2019a). Persistence of an endangered native duck, feral mallards, and multiple hybrid swarms across the main Hawaiian Islands. Molecular Ecology 28(24): 5203–5216.

Wells, C. P., P. Lavretsky, M. D. Sorenson, J. L. Peters, J. M. DaCosta, S. Turnbull, K. J. Uyehara, C. P. Malachowski, B. D. Dugger, J. M. Eadie, and A. Engilis Jr (2019b). Data for Persistence of an endangered native duck, feral mallards, and multiple hybrid swarms across the main Hawaiian Islands. Dryad dataset: 10.5061/dryad.ffbg79cqf

Whitehead, H. (2008). Analyzing animal societies: quantitative methods for vertebrate social analysis. University of Chicago Press.

Whitehead, H. (2009). SOCPROG programs: analysing animal social structures. Behavioral Ecology and Sociobiology 63: 765–778.

Zonana, D. M., J. M. Gee, E. S. Bridge, M. D. Breed, and D. F. Doak (2019). Assessing behavioral associations in a hybrid zone through social network analysis: complex assortative behaviors structure associations in a hybrid quail population. The American Naturalist 193(6): 852–865.

